# CEP41, a ciliopathy-linked centrosomal protein, regulates microtubule assembly and cell proliferation

**DOI:** 10.1101/2023.12.22.573068

**Authors:** Shweta Shyam Prassanawar, Tuhin Sarkar, Dulal Panda

## Abstract

Centrosomal proteins play pivotal roles in orchestrating microtubule dynamics, and their dysregulation leads to disorders, including cancer and ciliopathies. Understanding the multifaceted roles of centrosomal proteins is vital to comprehend their involvement in disease development. Here, we report novel cellular functions of CEP41, a centrosomal and ciliary protein implicated in Joubert syndrome. We show that CEP41 is an essential microtubule-associated protein with microtubule-stabilizing activity. Purified CEP41 binds to preformed microtubules, promotes microtubule nucleation, and suppresses microtubule disassembly. When overexpressed in cultured cells, CEP41 localizes to microtubules and promotes microtubule bundling. Conversely, shRNA-mediated knockdown of CEP41 disrupts the interphase microtubule network and delays microtubule reassembly, emphasizing its role in microtubule organization. Further, we demonstrate that CEP41’s association with microtubules relies on its conserved rhodanese homology domain (RHOD) and the N-terminal region. Interestingly, a disease-causing mutation in the RHOD domain impairs CEP41-microtubule interaction. Moreover, depletion of CEP41 inhibits cell proliferation and disrupts cell cycle progression, suggesting its potential involvement in cell cycle regulation. These insights into the cellular functions of CEP41 hold promise for unraveling the impact of its mutations in ciliopathies.

**Summary:** A novel role of CEP41 as a microtubule-associated protein that promotes microtubule assembly and regulates cell cycle progression could offer insights into the development of ciliopathies.

## Introduction

Several cellular processes essential for cell growth and survival rely on the dynamic remodeling of the microtubule network. In animal cells, the centrosome is the key organelle responsible for the controlled assembly and organization of microtubules (Conduit et al., 2015). Due to its pivotal role in microtubule regulation, centrosomes influence several microtubule-dependent cellular processes, including cell division, cell migration, organelle positioning, and maintaining cell shape and polarity. Particularly noteworthy is the centrosome’s role during mitosis, which ensures the formation of a dynamic bipolar spindle for precise chromosome segregation (Doxsey, 2001). Furthermore, the centrosome is a major signaling hub due to its indispensable role in ciliogenesis, involving several centrosomal proteins integral for proper cilia functioning (Bettencourt-Dias et al., 2011). Given these critical functions, defects in centrosome functioning adversely affect cell growth and development, resulting in a spectrum of diseases from cancer to developmental anomalies (Bettencourt-Dias et al., 2011; Nigg and Raff, 2009). Specifically, mutations in centrosomal proteins functioning at the cilium-centrosome interface have been linked to diverse genetic disorders broadly classified as ciliopathies (Vertii et al., 2015). These include developmental disorders such as Joubert syndrome, Bardet-Biedl syndrome, Meckel-Gruber syndrome, and orofaciodigital syndrome (Reiter and Leroux, 2017). Understanding the cellular functions of ciliopathy-associated centrosomal proteins is crucial for comprehending their role in disease development and devising targeted therapeutic strategies.

CEP41 is a centrosomal protein mutated in neurodevelopmental disorders like Joubert syndrome and autism (Korvatska et al., 2011; Lee et al., 2012; Patowary et al., 2019). CEP41 is essential for cilia assembly and indispensable in maintaining ciliary structure and motility by promoting ciliary localization of tubulin glutamylase (Lee et al., 2012). CEP41 has also been reported to be necessary for cilia reabsorption, with its involvement in activating the Aurora kinase A-dependent cilia disassembly pathway (Ki et al., 2020). Knockdown of CEP41 in zebrafish embryos severely hampers development due to disrupted ciliary functions and vascular impairment (Ki et al., 2020; Lee et al., 2012). Although CEP41’s ciliary functions have been well-studied, its role at the centrosome remains unexplored. In this regard, it is of significant interest to gain further information about the cellular functions of CEP41 to comprehend the impact of its mutations in ciliopathies.

Since CEP41 is associated with microtubule-based structures in cells (Gache et al., 2010; Lee et al., 2012), we aimed to explore its microtubule-associated functions. We provide evidence that CEP41 is a novel microtubule-associated protein (MAP) with microtubule-stabilizing activity. We used purified proteins to show that CEP41 binds to microtubules, promotes microtubule assembly, and stabilizes microtubules against disassembly. In cultured cells, CEP41 stabilizes microtubules and is essential for microtubule organization and cell proliferation. We also show that CEP41 interacts with the microtubules through its rhodanese-like domain and the coiled-coil motifs. Further, our study demonstrates CEP41’s involvement in mitosis, hinting at its yet unexplored role in cell cycle regulation. The findings advance our understanding of CEP41 as a microtubule-associated protein potentially involved in maintaining the microtubule network in cells.

## Results

### CEP41 interacts with tubulin and binds to microtubules

Based on previously reported localization of CEP41 at different microtubule-based structures, such as the centrosome, cilia, and basal bodies (Andersen et al., 2003; Jakobsen et al., 2011; Lee et al., 2012), we hypothesized that CEP41 could be a microtubule-interacting protein with microtubule-associated functions. We performed a co-immunoprecipitation assay to test whether CEP41 interacts with tubulin. CEP41 antibody, when incubated with the whole cell lysate of NIH3T3 cells, co-immunoprecipitated tubulin, indicating a physical interaction between the two proteins (Fig. 1A). Likewise, CEP41 was detected in the immunoprecipitants of α-tubulin, confirming that it is a novel binding partner of tubulin (Fig. 1B). As negative controls, rabbit IgG, mouse IgG, and Protein A agarose beads did not precipitate tubulin or CEP41. As positive controls, the presence of CEP41 and α-tubulin were detected in the immunoprecipitants of their respective antibodies. The positive interaction of CEP41 with tubulin prompted us to probe its interaction with microtubules further. We first checked if CEP41 is a part of the MAP family of proteins. MAPs are microtubule-binding proteins capable of modulating microtubule dynamics by imparting stabilizing or destabilizing effects (Bodakuntla et al., 2019). Brain tissue is a rich source of tubulin and MAPs, which can be fractionated by repeated cycles of tubulin polymerization and depolymerization, yielding tubulin rich with MAPs (Vallee, 1986). CEP41 was present in the crude brain tissue extract and MAP-rich fractions (Fig. 1C). Similarly, EB1, a plus-end microtubule-binding protein, was detected in the MAP-rich fractions. In contrast, Histone H3, a DNA-binding protein used as a negative control, did not fractionate with MAPs. The presence of CEP41 in both the first and second cycles of MAP-rich fractions indicates that CEP41 can bind and co-sediment with microtubules. It also suggests that CEP41 cycles with other MAPs like EB1 and can be a novel member of the microtubule-associated protein family.

**Fig 1.**
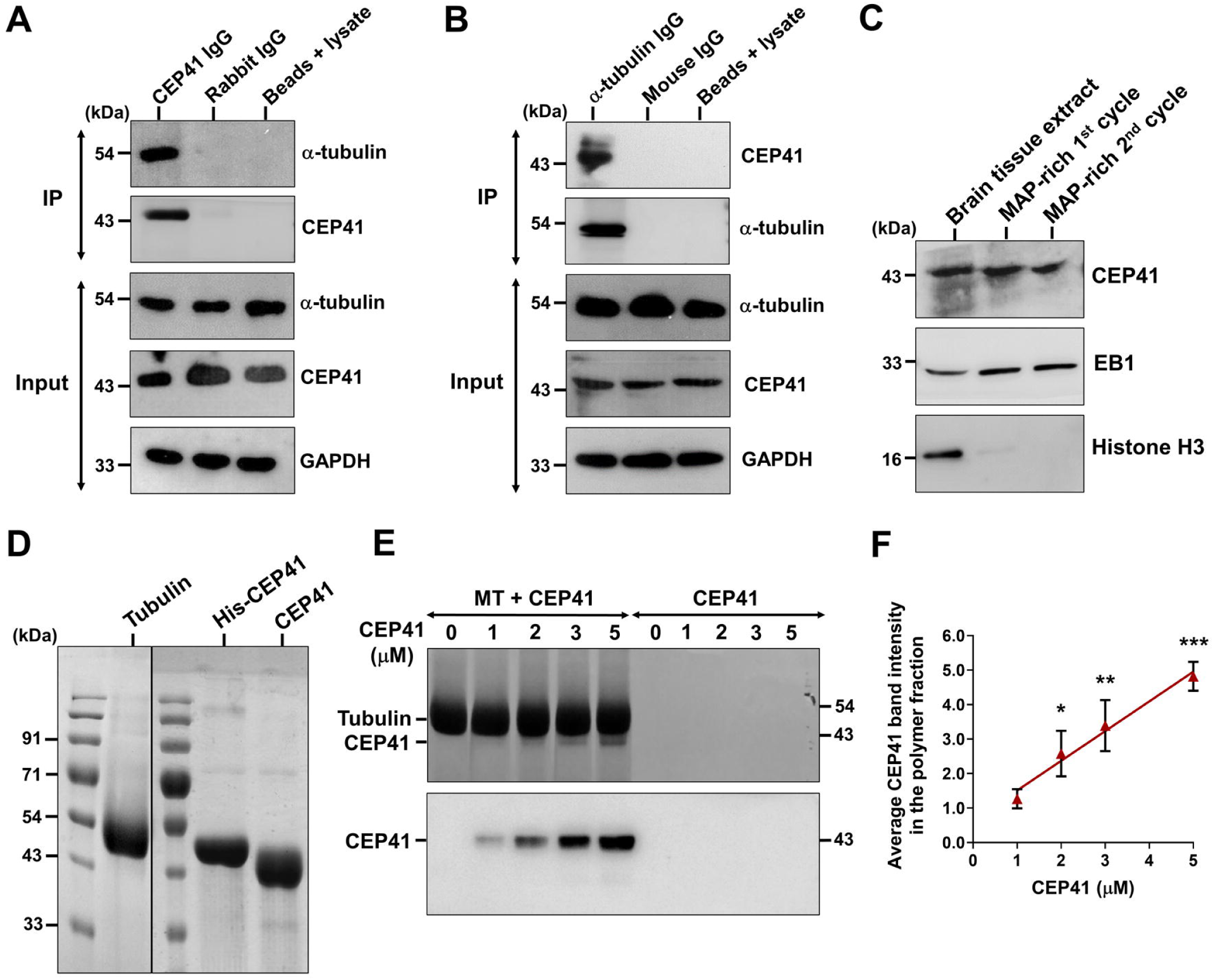
CEP41 interacts with tubulin and binds to microtubules. Co-immunoprecipitation of CEP41 and α-tubulin from NIH3T3 cells. The whole cell lysate was incubated with **(A)** CEP41 IgG or **(B)** α-tubulin IgG and immunoprecipitated using Protein A agarose beads. The inputs and eluates (IP) were immunoblotted with indicated antibodies. The experiments were performed twice, and representative blots are shown. **(C)** CEP41 was present in the MAP-rich tubulin fraction isolated from goat brain tissue. MAP-rich tubulin was isolated from a goat brain by two cycles of polymerization and depolymerization using 4 M glycerol. An equal concentration of fractions was loaded on SDS-PAGE gel and immunoblotted with CEP41 antibody. The experiment was performed twice, and a representative blot is shown. **(D)** Coomassie Brilliant Blue stained SDS-PAGE gel image of purified tubulin, his-CEP41, and his-tag cleaved CEP41. **(E)** A representative SDS-PAGE gel image (top) and immunoblot (bottom) of the co-sedimentation assay performed with CEP41 and preformed microtubules. Taxol-stabilized microtubules (3 μM) were incubated with indicated concentrations of CEP41 at 37°C. The polymers were sedimented by ultra-centrifugation, and an equal volume of the pellet fraction was resolved on SDS-PAGE gel and immunoblotted with anti-CEP41 antibody (bottom). **(F)** Band intensities of CEP41 in the pellet fraction were quantified using ImageJ. The experiment was performed three times, and error bars denote the standard deviation. Statistical significance was determined by Student’s *t*-test (\**P* < 0.05; \*\**P* < 0.01; \*\*\**P* < 0.001).

Next, we sought to establish a direct interaction between CEP41 and microtubules *in vitro*. To this end, we cloned and expressed histidine-tagged CEP41 in bacteria and affinity purified the protein (Fig. 1D) and validated it by immunoblotting with appropriate antibodies (Fig. S1A). Using purified proteins, we assessed the physical interaction of CEP41 with brain microtubules in a microtubule pelleting assay (Fig. 1E). Preformed taxol-stabilized microtubules were incubated with CEP41 and then sedimented by ultracentrifugation. CEP41 pelleted with the microtubules and was detected in the polymer fraction, whereas without microtubules, it did not sediment (Fig. 1E). The band intensity of the pelleted CEP41 increased significantly in a concentration-dependent manner (Fig. 1F), indicating that it was physically bound to microtubules and sedimented along with microtubules (Fig. 1F). To confirm that the sedimentation of CEP41 with microtubules was attributed to their interaction and not its aggregation, CEP41 was sedimented in similar conditions without microtubules and, as expected, remained in the soluble fraction (Fig. S1B).

### CEP41 promotes microtubule nucleation *in vitro*

The interaction of CEP41 with microtubules prompted us to check whether CEP41 regulates microtubule assembly. Spontaneous polymerization of microtubules *in vitro* occurs at higher tubulin concentrations as microtubule nucleation is a kinetically unfavorable process and requires the formation of stable tubulin oligomers (Roostalu and Surrey, 2017). Microtubule stabilizers, such as taxol and certain MAPs, enhance tubulin-tubulin interactions and promote microtubule assembly (Bre and Karsenti, 1990; Kumar, 1981; Tovey and Conduit, 2018). At first, we checked if CEP41 could induce spontaneous tubulin polymerization without any polymerization promoter. Interestingly, CEP41 polymerized tubulin in the presence of GTP, as observed in a turbidity-based assay (Fig. 2A). CEP41-induced tubulin polymerization showed a classic sigmoidal curve with an initial lag phase followed by an elongation phase that saturated over time (Fig. 2A). With increasing CEP41 concentrations, the lag time decreased, and the microtubule mass increased. Tubulin alone at a similar concentration did not polymerize. Also, bovine serum albumin and alkaline phosphatase as negative controls did not induce microtubule assembly. Compared to the control, where tubulin did not polymerize, in the presence of 1 μM CEP41, there was a 10-fold increase in the assembly rate (Fig. 2B). Furthermore, the rate increased by 13, 39, and 59-fold in the presence of 2, 3, and 5 μM CEP41, respectively (Fig. 2B). Next, we sedimented the microtubules formed in the presence of CEP41 and observed a distinct increase in the polymer mass with increasing concentrations of CEP41 (Fig. 2C). In the absence of CEP41, tubulin did not polymerize, as evident by the less sedimented mass. The band intensities of the pelleted microtubules increased significantly with increasing CEP41. Likewise, increasing concentrations of CEP41 reduced the mass of unpolymerized soluble tubulin, indicating tubulin heterodimers polymerizing into microtubules (Fig. 2C). The polymer/soluble ratio increased by 1.8, 2.3, 3, and 3.9-fold in the presence of 1, 2, 3, and 5 μM CEP41, respectively (Fig. 2D).

**Fig 2.**
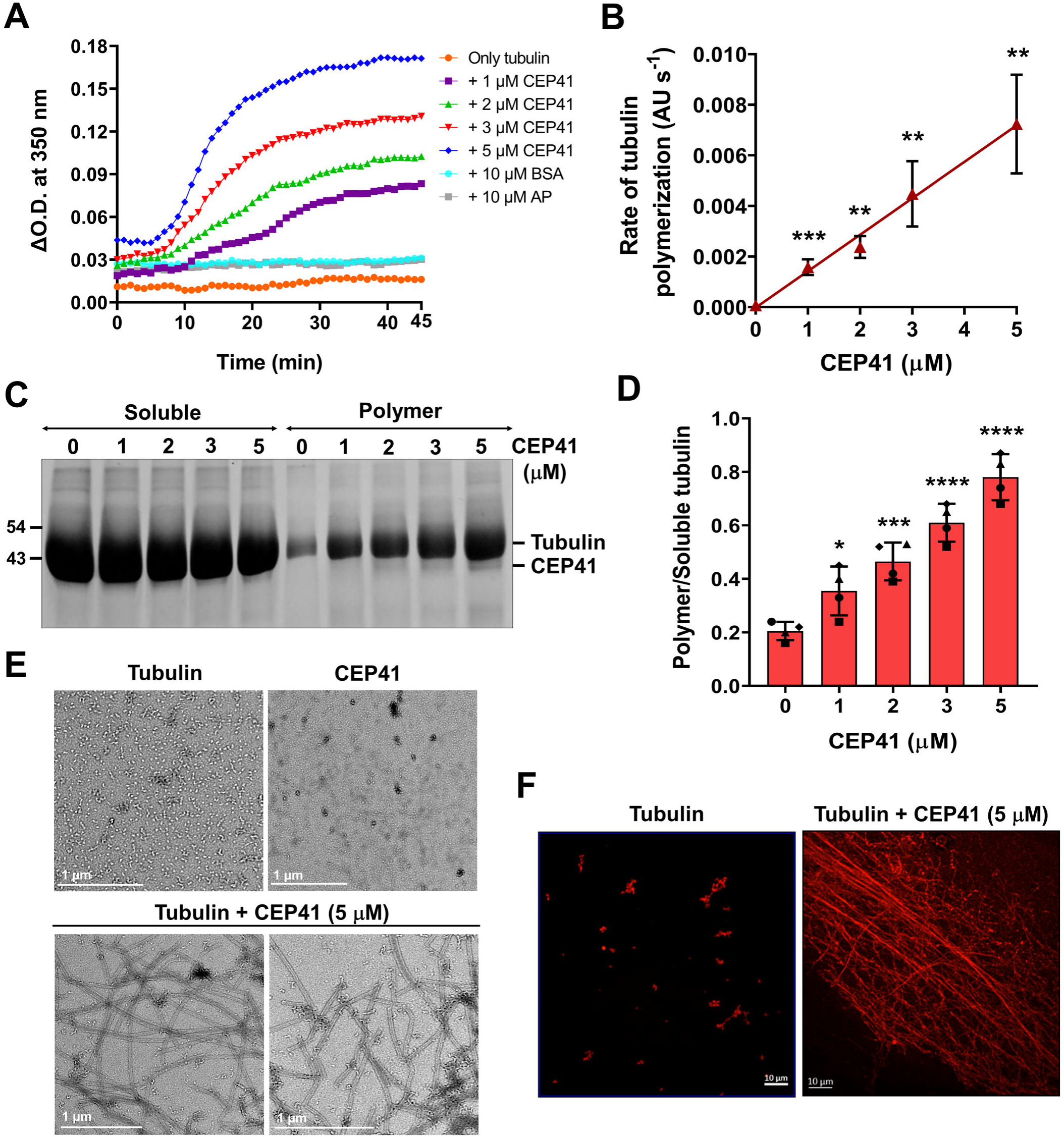
CEP41 promoted tubulin assembly in the presence of GTP. **(A)** *In vitro* polymerization of tubulin (30 μM) without (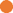) or with 1 (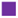), 2 (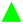), 3 (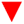), and 5 (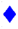) μM CEP41 and 1 mM GTP was monitored at 37°C. Bovine serum albumin (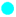) and alkaline phosphatase (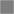) were negative controls. The experiment was performed thrice, and a representative plot is shown. **(B)** The rate of tubulin polymerization was calculated from the slope of the elongation phase (\*\**P* < 0.01; \*\*\**P* < 0.001). Error bars represent the standard deviation from three independent sets of experiments. **(C)** Tubulin (30 μM) was polymerized in the absence or presence of 1, 2, 3, and 5 μM CEP41 for 45 min at 37°C, and the soluble and polymer fractions were separated by ultra-centrifugation. An equal volume of the fractions was resolved on SDS-PAGE gel; a representative gel image is shown. **(D)** Band intensities of tubulin in the soluble and the polymer fractions were quantified using ImageJ, and the polymer/soluble ratio was determined (\**P* < 0.05; \*\*\**P* < 0.001; \*\*\*\**P* < 0.0001). Error bars represent the standard deviation from four independent sets of experiments. Statistical significance was determined by Student’s *t*-test. **(E)** Transmission electron microscopy images of microtubules formed in the presence of CEP41. Tubulin (30 μM) was polymerized in the presence of 5 μM CEP41 for 45 min at 37°C. The scale bar is 1 μm. **(F)** Tubulin was polymerized in the same manner mentioned above. The reaction mixture was loaded on poly-L-lysine coated coverslips, immunostained with α-tubulin antibody, and imaged in a spinning-disk confocal microscope at 63x magnification. The scale bar is 10 μm. The experiments were performed three times, and representative images are shown.

We also visualized the microtubules formed in the presence of CEP41 using transmission electron microscopy (Fig. 2E). Polymers were not observed in the absence of CEP41 when tubulin was incubated only with GTP (Fig. 2E, top left panel). Similarly, CEP41 without tubulin showed no polymer formation (Fig. 2E, top right panel). In the presence of CEP41, microtubules were seen with a dense mesh-like network of polymers in most fields (Fig. 2E, bottom panels). We also visualized the CEP41-induced tubulin polymers by confocal microscopy by immunostaining the microtubules using α-tubulin antibody (Fig. 2F). As observed previously, here, too, in the absence of CEP41, there were no polymers and only small tubulin aggregates (Fig. 2F, left panel). In the presence of CEP41, long filaments in a mesh-like network were seen along with thick microtubule bundles (Fig. 2F, right panel). The results demonstrate that CEP41 can promote microtubule nucleation and polymerization *in vitro*.

### CEP41 enhances the rate of taxol-induced microtubule assembly

Next, we examined the effect of CEP41 on microtubule assembly in the presence of taxol, which promotes tubulin polymerization at lower tubulin concentrations. CEP41 promoted tubulin assembly in the presence of taxol (Fig. 3A). Compared to the control, the extent of polymerization increased by 1.3, 1.4, 1.5, and 1.7-fold in the presence of 1, 2, 3, and 5 μM CEP41, respectively, as evident by an increase in light scattering. Also, CEP41 significantly increased the rate of tubulin polymerization, resulting in faster and more polymer formation (Fig. 3B). In comparison to the rate of polymerization in the presence of taxol, the rate increased by 1.4, 1.8, 2.1, and 2.3-fold in the presence of 1, 2, 3, and 5 μM CEP41, respectively, suggesting that CEP41 may assist in the nucleation phase of microtubule assembly. Similarly, in a sedimentation assay, compared to only taxol, more polymeric tubulin was sedimented in the presence of CEP41 (Fig. 3C). The band intensity of pelleted microtubules increased significantly with increasing CEP41 concentration. The polymer/soluble ratio increased by 1.3, 1.4, 1.7, and 1.9-fold in the presence of 1, 2, 3, and 5 µM CEP41, respectively (Fig. 3D). The data suggest that CEP41 is a promoter of microtubule assembly.

**Fig 3.**
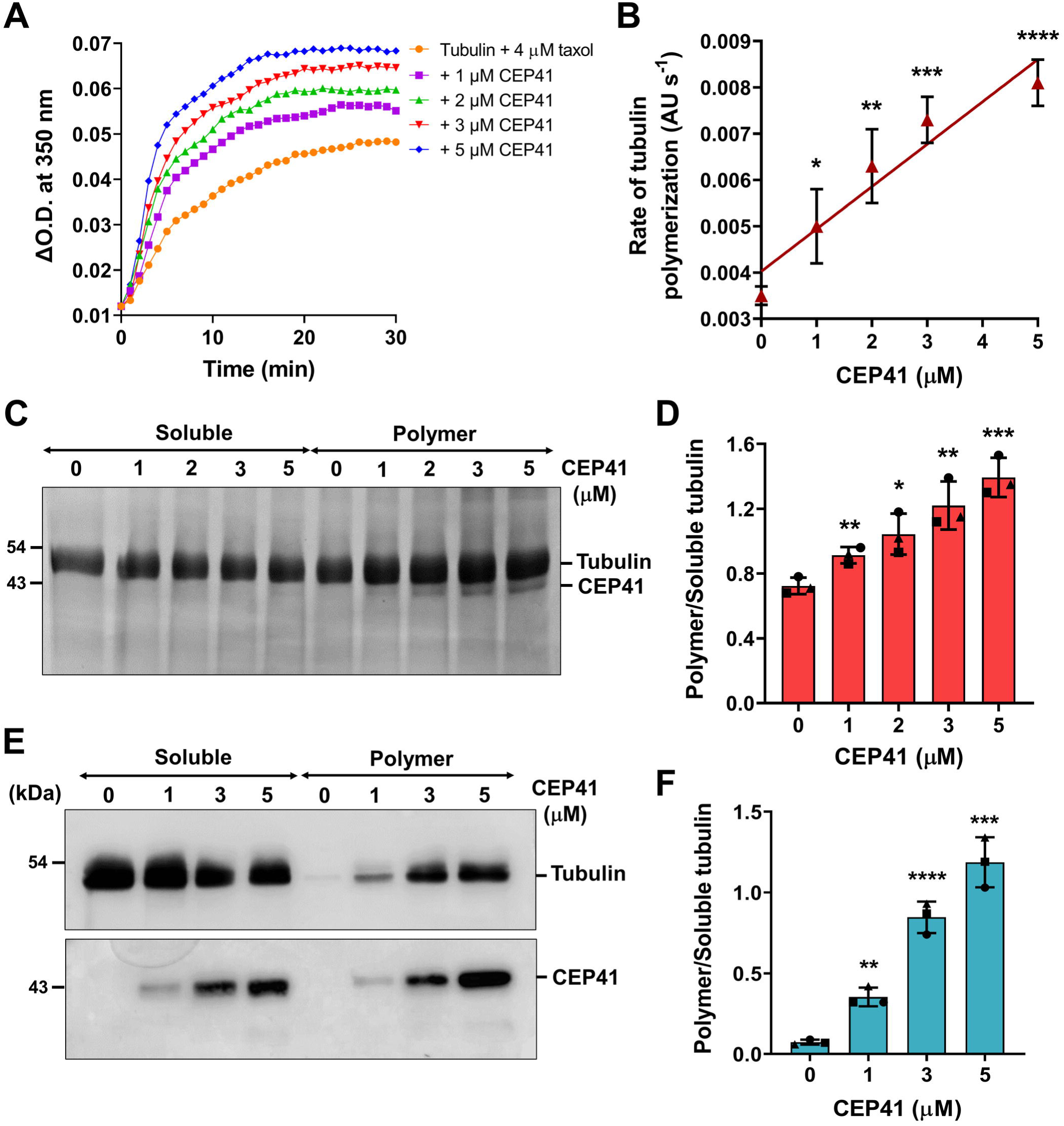
CEP41 enhanced the rate of taxol-induced tubulin polymerization and prevented dilution-induced microtubule disassembly. **(A)** The effect of CEP41 on taxol-induced tubulin polymerization was monitored at 37°C. Tubulin (15 μM) was incubated with different concentrations of CEP41 (0 (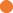), 1 (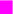), 2 (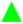), 3 (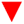), 5 (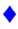) μM) followed by taxol (4 μM) and GTP (1 mM). The experiment was performed thrice, and a representative plot is shown. **(B)** The rate of tubulin polymerization was calculated from the slope of the elongation phase (log phase of the polymerization curve in **A** (\**P* < 0.05; \*\**P* < 0.01; \*\*\**P* < 0.001; \*\*\*\**P* < 0.0001). Error bars represent the standard deviation from three independent sets of experiments. **(C)** Tubulin (15 μM) was polymerized in the presence of taxol (4 μM), and different concentrations (0, 1, 2, 3, and 5 μM) of CEP41 for 30 min at 37°C and the soluble and polymer fractions were separated by ultra-centrifugation. An equal volume of soluble and polymer fractions was resolved on SDS-PAGE gel, and a representative gel image is shown. **(D)** Band intensities of tubulin in the soluble and the polymer fraction were quantified using ImageJ, and the polymer/soluble ratio was calculated (\**P* < 0.05; \*\**P* < 0.01; \*\*\**P* < 0.001). Error bars represent the standard deviation from three independent sets of experiments. **(E)** Representative immunoblot of dilution-induced disassembly assay performed with CEP41 and preformed microtubules. Microtubules were polymerized in the presence of DMSO and diluted 10-fold, followed by incubation with indicated concentrations of CEP41 at 37°C. The polymers were sedimented by ultra-centrifugation, and equal volumes of the supernatant and pellet fractions were resolved on SDS-PAGE gel and immunoblotted with anti-α-tubulin (top) and anti-CEP41 antibody (bottom). **(F)** Band intensities of tubulin in the soluble and the polymer fractions were quantified using ImageJ, and the polymer/soluble ratio was calculated. The experiment was performed three times, and error bars denote the standard deviation. Statistical significance was determined by Student’s *t*-test (\*\**P* < 0.01; \*\*\**P* < 0.001; \*\*\*\**P* < 0.0001).

### CEP41 prevents dilution-induced disassembly of microtubules *in vitro*

Furthermore, we assessed the effect of CEP41 on dilution-induced microtubule disassembly. Microtubules undergo depolymerization upon dilution, as the concentration of tubulin subunits falls below the critical concentration required to maintain the assembly rate (Karr and Purich, 1979). We polymerized microtubules in the presence of DMSO and then diluted them 10-fold in both the absence and presence of CEP41. Microtubules resisted depolymerization when incubated with CEP41, as evidenced by more polymer mass sedimented compared to the control (Fig. 3E). In the presence of 1, 3, and 5 μM CEP41, the polymer/soluble tubulin ratio increased by 5, 10, and 16-fold, respectively, suggesting inhibition of microtubule depolymerization (Fig. 3F). Additionally, the presence of CEP41 in the polymer fractions complements the earlier microtubule-binding assay result, indicating that CEP41 interacts with the microtubules and sediments along with them (Fig. 3E). Together, the results suggest that CEP41 binds to microtubules and imparts a stabilizing effect, promoting assembly and preventing disassembly.

### Exogenously expressed CEP41 colocalizes with interphase microtubules and associates with the spindle poles and midbody during mitosis

Previous studies have reported the localization of CEP41 at the cilia and basal bodies and established its essential role in cilia functioning (Ki et al., 2020; Lee et al., 2012). We wanted to investigate the cilia-unrelated functions of CEP41 and characterize its localization during cell division. Since the polyclonal antibody available for CEP41 was unsuitable for immunofluorescence analysis, we exogenously expressed GFP-tagged CEP41 in human cervical cancer (HeLa) and mouse fibroblast (NIH3T3) cells. In HeLa cells, CEP41-GFP, when expressed in low amounts, showed centrosomal localization, whereas, in high-expressing cells, it localized to microtubules and showed a dense fiber-like network spread throughout the cell (Fig. 4A; Fig. S2A,B). Similar CEP41 localization was observed in NIH3T3 cells as well (Fig. S2C). A distinct overlap between CEP41-GFP and α-tubulin fluorescence was visible, indicating colocalization (Fig. 4A). In contrast, GFP did not show such colocalization and exhibited diffused staining in the cytoplasm and nucleus in both cell lines (Fig. 4A; Fig. S2). CEP41-GFP localizing on microtubules further strengthens our findings that it binds and interacts with microtubules in cells. To measure the extent of colocalization, we determined Pearson’s correlation coefficient (*r*), which measures the level of correlation between the two fluorophores. An *r* value close to 1 indicates a strong correlation, whereas a value close to 0 or negative indicates no correlation. The coefficient value *r* for CEP41 and microtubules was estimated to be 0.8 ± 0.01, indicating a strong correlation between the fluorophores (Fig. 4B). The *r* value for GFP alone and microtubules was −0.25 ± 0.05, showing no correlation.

**Fig 4.**
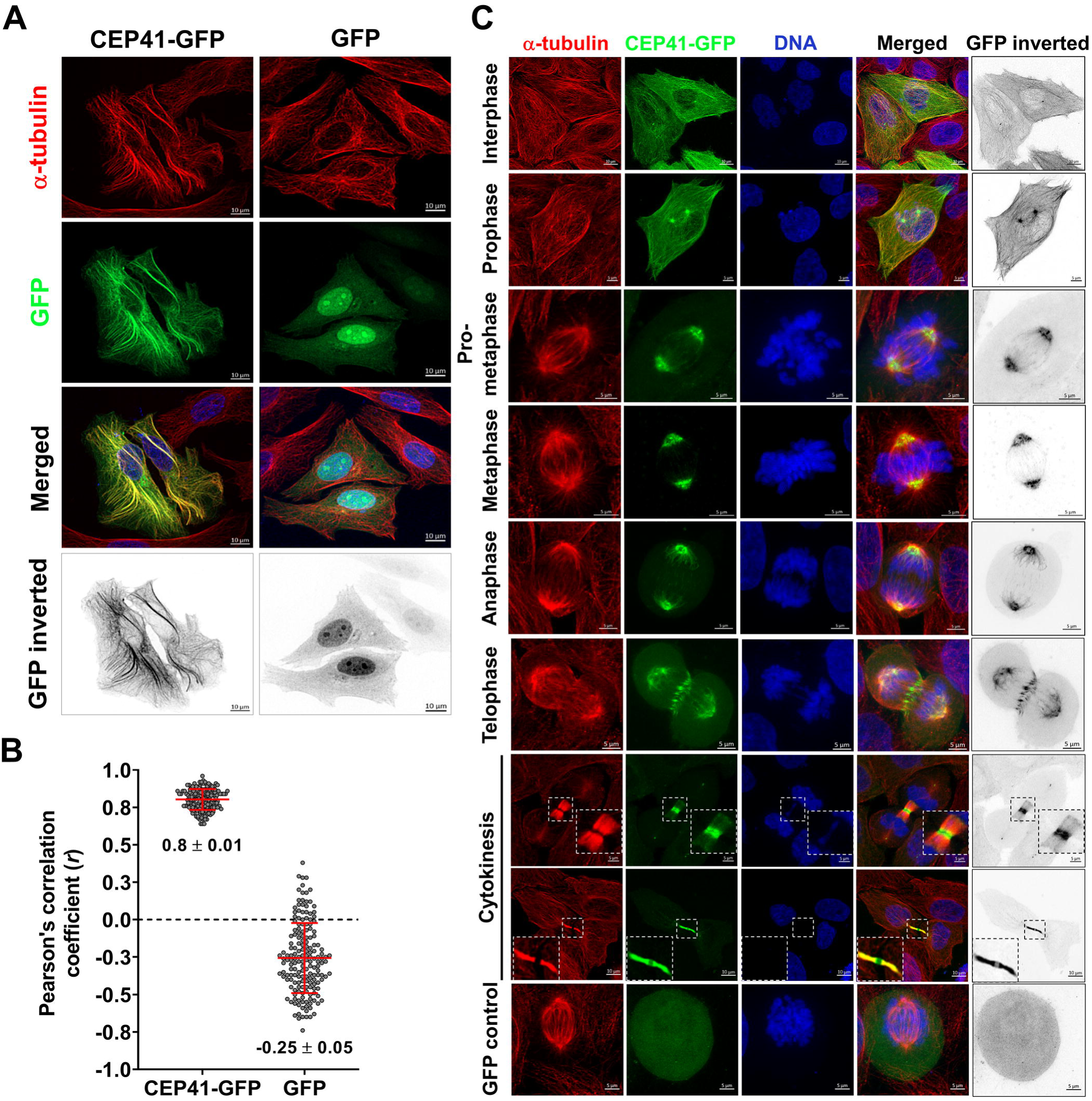
Exogeneously expressed CEP41 colocalized with microtubules and associated with spindle poles and midbody in HeLa cells. **(A)** Fluorescence microscopy images of HeLa cells transfected with CEP41-GFP or GFP constructs. After 24 h of transfection, cells were fixed with 3.7% paraformaldehyde and immunostained for α-tubulin (red). DNA was stained with Hoechst (blue). The scale bar is 10 μm. **(B)** Pearson’s correlation coefficient (*r*) of GFP-expressing cells was determined using the Coloc 2 plugin in ImageJ. Data represent the mean ± s.d. of three independent experiments with 170 cells analyzed in each group. **(C)** Localization of CEP41-GFP at different stages of the cell cycle. A double thymidine block was used to synchronize HeLa cells transfected with either CEP41-GFP or GFP constructs to enrich mitotic cells. The cells were fixed and stained for α-tubulin (red) and DNA (blue). Images were captured in a laser scanning confocal microscope at 63x magnification. Scale bar: 5 μm and the insets show magnification of the midbody. The bottom last panel is for GFP control showing no GFP localization at the spindle poles.

We next determined the localization of CEP41 during different cell cycle stages. In HeLa cells, CEP41-GFP localized to the spindle poles and microtubule-based structures throughout mitosis (Fig. 4C). In interphase and prophase, it localized to the centrosome and cytoplasmic microtubules and, upon nuclear envelope breakdown, was prominently seen at both the spindle poles and spindle microtubules, specifically near the poles. During telophase, CEP41 accumulated at the intermediate region of the cleavage furrow and, throughout cytokinesis, remained associated with the midbody. Similar localization of CEP41 at the midbody was observed in NIH3T3 cells (Fig. S2D), and GFP as negative control did not localize to the mitotic spindle (Fig. 4C). Our result is the first evidence to show the involvement of CEP41 during cell division, suggesting its potential functions in mitosis and cytokinesis.

### CEP41 interacts with microtubules through its rhodanese-like domain and the N-terminal region

As CEP41 lacked any known microtubule-binding domain, we sought to identify its microtubule-interacting region. Prediction of its secondary and tertiary structure revealed that CEP41 comprises two coiled-coil motifs at the N-terminal region and a rhodanese-like domain (RHOD) at the intermediate region. In contrast, the C-terminal segment of the protein appeared to be primarily disordered (Fig. 5A; Fig. S3A,B). To investigate this further, we generated several deletion constructs of CEP41 fused with GFP at the C-terminal and expressed the truncated proteins in HeLa cells through transient transfection (Fig. 5B,C; Fig. S4). In the case of the N-terminal deletion construct (CEP41^ΔN^), the protein did not localize to microtubules and instead exhibited diffused localization in the cytoplasm, similar to the GFP control (Fig. 5C). For the construct with the rhodanese domain deleted (CEP41^ΔRHOD^), we noticed the formation of substantial aggregates in the cytoplasm, likely stemming from protein misfolding following the deletion of a conserved domain. Interestingly, the CEP41 construct with the C-terminal region deleted displayed a localization pattern similar to that of the full-length protein and exhibited colocalization with microtubules. This suggests that the C-terminal disordered region is dispensable for CEP41-microtubule interaction. Moreover, when only the RHOD domain was expressed without the N and C-terminal regions, the protein displayed a diffused distribution in the cytoplasm (Fig. 5C). These observations suggest that the coiled-coil motifs and the RHOD domain are essential for CEP41’s interaction with microtubules. In dividing cells, all the truncated mutants localized to the spindle poles; however, only the CEP41 construct with the C-terminal region deleted localized to the midbody (Fig. 5D; Fig. S4B,C). This suggests that the RHOD domain plays a role in CEP41-microtubule interaction rather than centrosomal localization. Remarkably, the coiled-coil motifs and the RHOD domain are highly conserved among CEP41 orthologues, implying their essential role in CEP41’s cellular functions (Fig. S3C). When we tested some of the known disease-causing mutations of CEP41 for their ability to interact with microtubules, one mutant with a mutation in the RHOD domain, R179H, did not localize to microtubules, indicating impaired interaction (Figure 5C). Conversely, other tested mutants, with mutations located in the coiled-coil motifs (M36T and Q89E) and the RHOD domain (P206A), exhibited microtubule localization and bundling similar to wild-type CEP41 (Fig. S5). This underscores the significance of the RHOD domain and specifically of the R179 residue in mediating CEP41-microtubule interaction.

**Fig 5.**
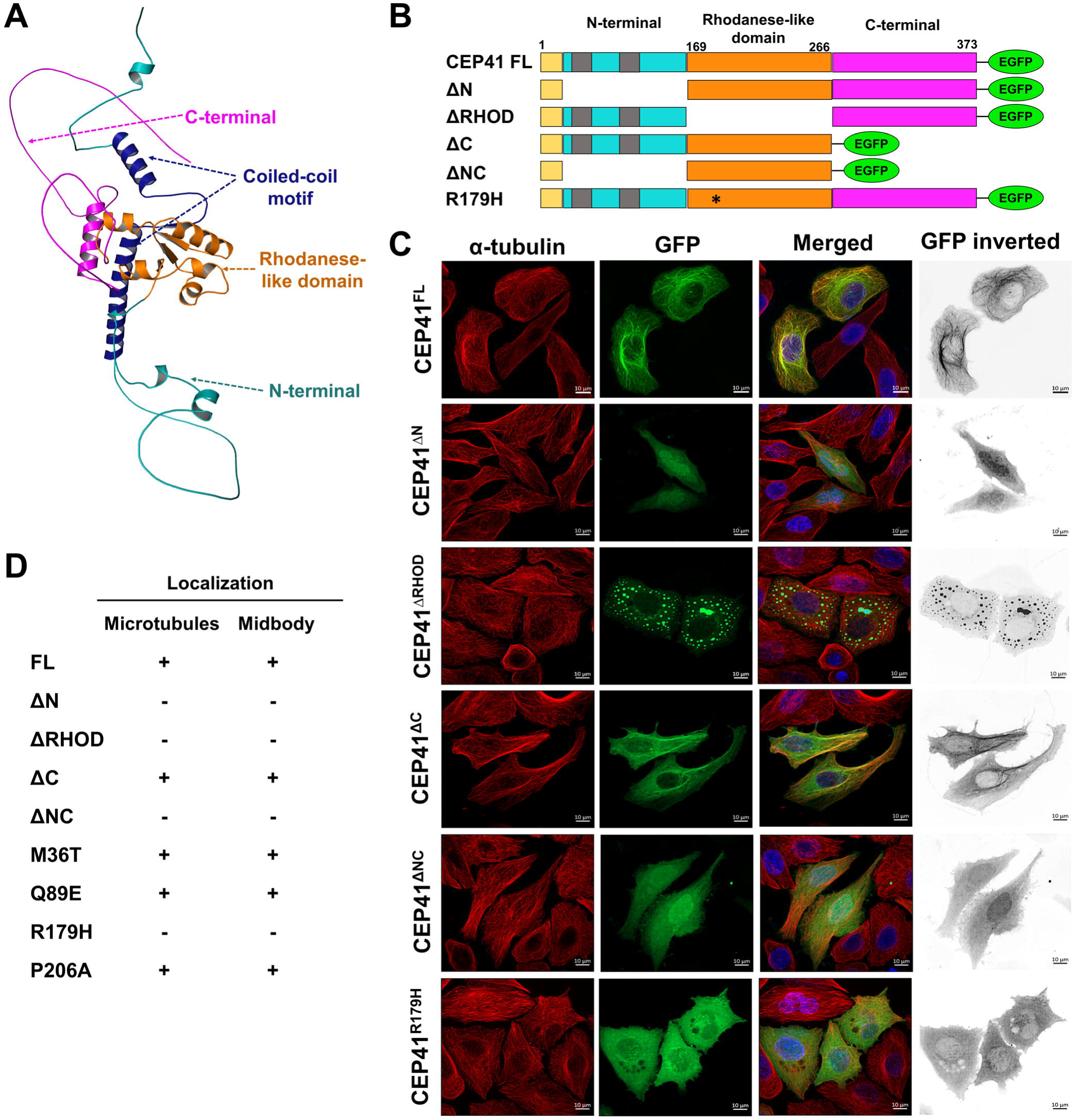
CEP41 interacted with microtubules via its RHOD domain and the N-terminal region. **(A)** AlphaFold-predicted structure of CEP41. Color code, N-terminal region: cyan, coiled-coil motifs: dark blue, RHOD domain: orange, C-terminal region: pink. The image was created using PyMoL. (B) Deletion and mutated constructs of CEP41. The yellow box indicates the predicted nuclear exit sequence, and the grey boxes indicate the predicted coiled-coil motifs. **(C)** Fluorescence microscopy images of HeLa cells transfected with GFP-tagged full-length and truncated CEP41 constructs (green). Transfected cells were immunostained for α-tubulin (red) 24 h post-transfection. DNA was stained with Hoechst (blue). Images were captured in a spinning-disk confocal microscope at 63x magnification. Scale bar: 10 μm. **(D)** Summarized results of the localization of GFP-tagged truncated and mutated CEP41. The experiment was performed twice, with 100 cells analyzed in each group.

### Overexpression of CEP41 induces the formation of stable microtubule bundles

We also observed noticeable microtubule bundling in HeLa cells overexpressing CEP41 (Fig. 6A). Most CEP41-overexpressing cells exhibited thick bundles of straight microtubules (Fig. 6A, i), while in some cases, the microtubules were concentrated in a circular ring around the nucleus (Fig. 6A, ii). No significant microtubule bundling was observed in GFP control cells, as they showed a well-spread interphase arrangement of microtubules. We counted the number of cells with prominent microtubule bundles, and 72 ± 12% of the cells expressing CEP41-GFP showed microtubule bundling (Fig. 6B). In contrast, only 15 ± 2% of GFP-expressing cells showed distinct bundles. Due to significant bundling, we expected a concomitant increase in the microtubule intensity, so we quantified the mean microtubule intensity per cell. The fluorescence intensity increased by 42 ± 4% in CEP41-overexpressing cells compared to non-transfected control cells, with no significant change in GFP-control cells (Fig. 6C). Next, we assessed the stability of these microtubule bundles. CEP41-induced microtubule bundles were positively stained with acetylated tubulin antibody (Fig. 6D). Acetylation of the Lys40 residue is a post-translational modification of α-tubulin and a marker for stable, long-lived microtubules (Fig. 6D) (Piperno et al., 1987). Non-transfected control and GFP-control cells showed no significant microtubule bundling and a basal level of acetylated tubulin staining, whereas the acetylated tubulin intensity increased by 81 ± 18% in CEP41-overexpressing cells (Fig. 6D). Further, we examined the stability of CEP41-induced microtubule bundles against the depolymerizing action of nocodazole. The interphase microtubule network depolymerized in control and GFP-expressing cells (Fig. 6E). However, in CEP41-GFP expressing cells, prominent microtubule bundles were visible, which were resistant against nocodazole-induced depolymerization. We quantified the mean microtubule intensity per cell, which was increased by 75 ± 12% in CEP41-overexpressing cells compared to the control (Fig. 6E). The formation of stable microtubule bundles and localization of CEP41 to these bundles hints toward a possible role of CEP41 in microtubule cross-linking. Together with the *in vitro* findings, the results suggest that CEP41 is a microtubule-stabilizing protein involved in microtubule nucleation and organization. Given that CEP41 is known to participate in ciliary glutamylation, we examined polyglutamylated tubulin levels in CEP41-overexpressing cells (Fig. S6). However, compared to non-transfected control cells, CEP41-overexpressing cells did not exhibit a significant increase in polyglutamylated tubulin intensity. This implies that while CEP41 plays a role in ciliary tubulin glutamylation, it does not significantly influence cytoplasmic tubulin glutamylation.

**Fig 6.**
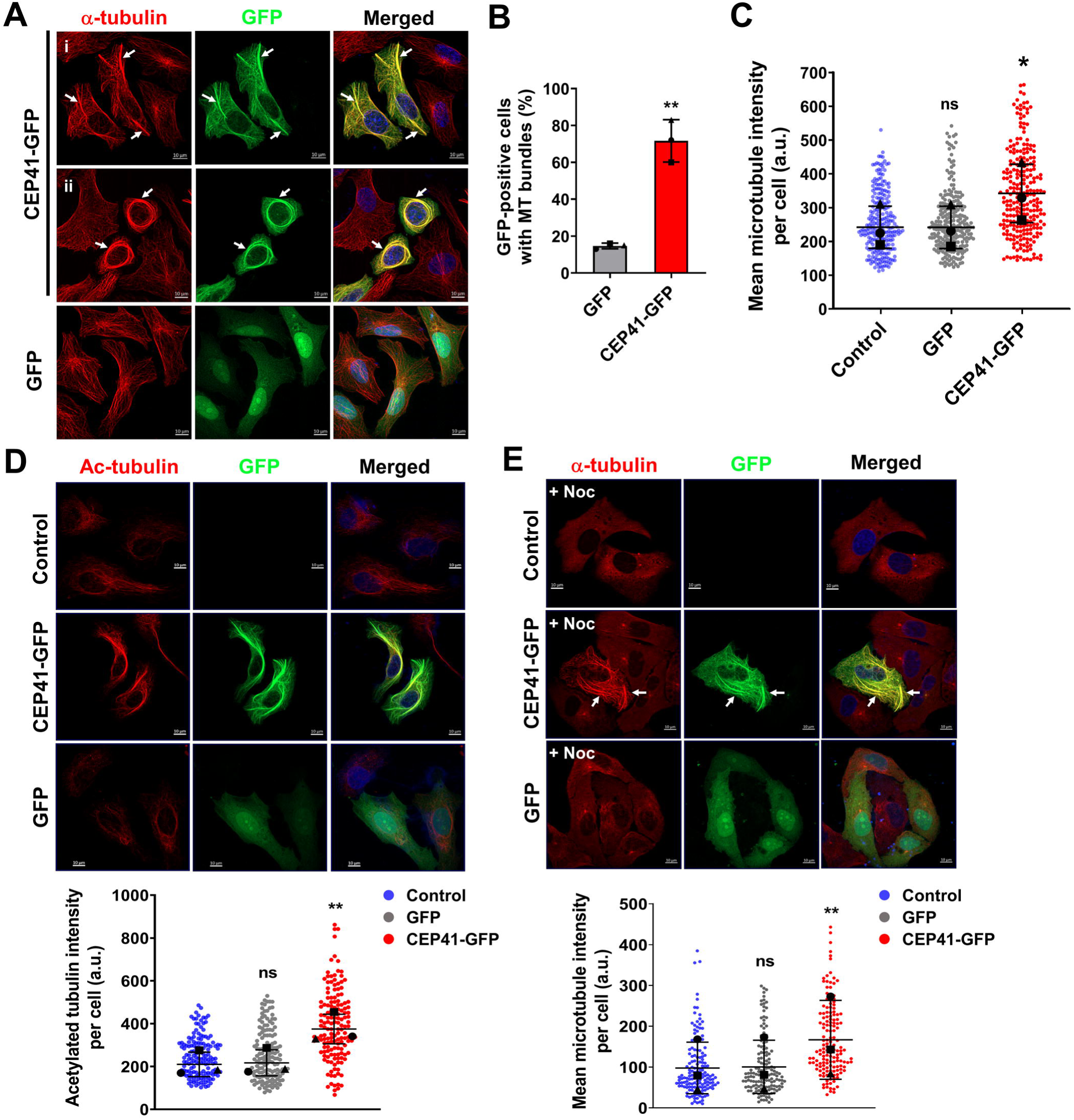
Overexpression of CEP41 induced the formation of stable microtubule bundles. **(A)** Fluorescence microscopy images of HeLa cells transfected with CEP41-GFP or GFP constructs and immunostained for α-tubulin (red). DNA was stained with Hoechst (blue). The scale bar is 10 μm, and white arrows point to microtubule bundles. Panels, i and ii, show different forms of microtubule bundles formed in HeLa cells upon CEP41 overexpression. **(B)** Percentage of cells with microtubule bundles plotted for GFP and CEP41-GFP expressing cells. The data are an average of three independent experiments with 300 cells counted in each case (ns *P* > 0.05; \*\**P* < 0.01). **(C)** Microtubule intensity of 250 cells in each case was quantified using ImageJ (ns *P* > 0.05; \**P* < 0.05). Data represent the mean ± s.d. of three independent experiments. **(D)** CEP41 overexpression enhanced the level of tubulin acetylation. HeLa cells transfected with CEP41-GFP and GFP constructs were stained for acetylated tubulin (red) and DNA (blue), scale bar: 10 μm. The intensity of acetylated tubulin per cell was quantified for 150 cells in each case (ns *P* > 0.05; \*\**P* < 0.01). **(E)** Exogenous expression of CEP41 stabilized microtubules against nocodazole-induced depolymerization. HeLa cells transfected with GFP, or CEP41-GFP constructs, were treated with nocodazole (500 nM) for 1 h and stained for α-tubulin (red) and DNA (blue), scale bar: 10 μm. The white arrows mark microtubule bundles. Microtubule intensity of 150 cells in each case was quantified using ImageJ (ns *P* > 0.05; \*\**P* < 0.01). The experiments were performed three times, and error bars denote standard deviation. Statistical significance was determined by Student’s *t*-test.

### Depletion of CEP41 depolymerizes the interphase microtubule network and reduces the extent of microtubule reassembly

We further elucidated the role of CEP41 in microtubule regulation via loss of function experiments where we depleted the endogenous levels of CEP41 using shRNA. Given that CEP41 localization was similar in HeLa and NIH3T3 cells, we chose the NIH3T3 cell line for the depletion experiments as it is a non-cancerous fibroblast cell line capable of undergoing ciliogenesis. CEP41 is an essential ciliary protein reported to be crucial for ciliogenesis and cilia functioning and is expressed in more amounts in NIH3T3 than in HeLa cells (Graser et al., 2007; Ki et al., 2020; Lee et al., 2012). Among the two shRNAs targeting CEP41, shRNA-2 showed a considerable reduction in CEP41 levels (Fig. S7A). Immunoblotting of NIH3T3 cell lysate, 48 h post-transfection with CEP41 shRNA-2, showed a 74 ± 2% reduction in the protein level compared to untransfected and scrambled control (Fig. 7A). Further, immunofluorescence analysis of CEP41-depleted cells revealed that CEP41 depletion depolymerized microtubules and disrupted the interphase microtubule network (Fig. 7B). Control and scrambled control cells exhibited a well-spread microtubule network with thin hair-like filaments throughout the cell. However, CEP41-depleted cells lacked a well-spread microtubule network and showed fewer microtubules (Fig. 7B). We quantified the fluorescence intensity of microtubules per cell, which was reduced by 47 ± 1.2% in CEP41-depleted cells compared to the control cells (Fig. 7C). CEP41-depleted cells also exhibited a drastic change in cell morphology as they were significantly smaller than control cells. The cytoplasmic area of CEP41-depleted cells was reduced by 55 ± 7% compared to the control cells (Fig. 7C). Despite observing a significant decrease in microtubule intensity in CEP41-depleted cells, the total levels of tubulin remained unchanged when compared to control and scrambled control cells (Fig S7B,C), indicating that the reduction observed in immunofluorescence images corresponds to decreased levels of polymerized tubulin. Furthermore, CEP41 depletion did not affect polyglutamylated tubulin levels either (Fig. S7B,C). We also checked the effect of CEP41 depletion on the actin network of NIH3T3 cells, which exhibited no significant difference compared to scrambled control cells (Fig. S7B-E), further ensuring that the observed phenotype is a result of the effect of CEP41 depletion on microtubules.

**Fig 7.**
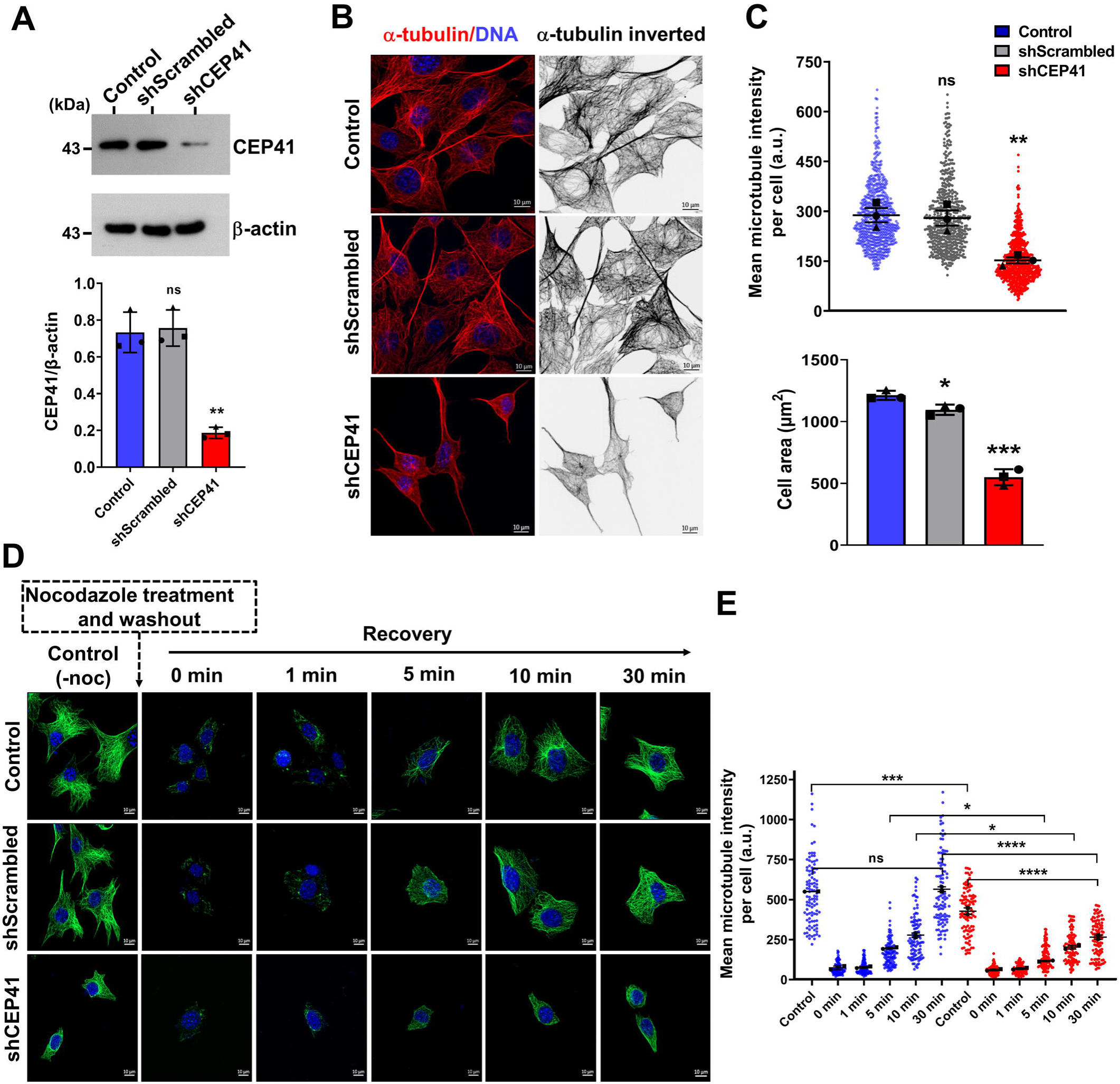
CEP41 depletion disrupted the interphase microtubule organization and reduced the extent of microtubule reassembly in NIH3T3 cells. **(A)** Representative immunoblot showing CEP41 levels in NIH3T3 cells transfected with scrambled or CEP41 shRNA. β-actin was used as a loading control. Band intensities were quantified using ImageJ, and the CEP41/β-actin intensity ratios were plotted (ns *P* > 0.05; \*\**P* < 0.01). Error bars represent the standard deviation from three independent sets of experiments. **(B)** Fluorescence microscopy images of NIH3T3 cells transfected with scrambled or CEP41 shRNA. Transfected cells were fixed and stained for α-tubulin (red) and DNA (blue). Images were captured in a spinning-disk confocal microscope at 63x magnification. Scale bar: 10 μm. **(C)** Microtubule intensity and cell area of 250 cells in each case were quantified using ImageJ (ns *P* > 0.05; \**P* < 0.05; \*\**P* < 0.01; \*\*\**P* < 0.001). Data represent the mean ± s.d. of three independent sets. Statistical significance was determined by Student’s *t*-test. **(D)** Microtubule reassembly assay. NIH3T3 cells transfected with scrambled or CEP41 shRNA were treated with nocodazole (500 nM) for 1 h to disassemble interphase microtubules. Post-treatment, nocodazole was washed out, and microtubule reassembly kinetics were monitored by incubating cells at 37°C. At indicated time points, the soluble fraction was extracted, and cells were fixed and stained for α-tubulin (green) and DNA (blue). Images were captured in a spinning-disk confocal microscope at 63x magnification. Scale bar: 10 μm. **(E)** Microtubule intensity of 100 cells in each case were quantified using ImageJ (ns *P* > 0.05; \**P* < 0.05; \*\*\**P* < 0.001; \*\*\*\**P* < 0.0001). Data represent the mean ± s.e.m. of two independent experiments. Statistical significance was determined by one-way ANOVA test.

To assess CEP41’s role in microtubule nucleation, we performed a nocodazole washout assay and monitored microtubule reassembly. NIH3T3 cells, either untransfected or transfected with scrambled or CEP41 shRNA, were treated with nocodazole for 1 h to disassemble interphase microtubules. Subsequently, nocodazole was washed out, and microtubule reassembly kinetics were monitored by incubating cells at 37°C for various time intervals (Fig. 7D). Before nocodazole treatment, CEP41-depleted cells exhibited reduced cell size and disrupted microtubule network, with mean microtubule intensity decreasing by 22 ± 4% compared to control (Fig. 7E). Upon nocodazole treatment, the interphase microtubule network depolymerized, and the microtubule intensity decreased by 87% in all groups. During recovery periods of 1, 5, 10, and 30 min, control cells showed a gradual increase in microtubule intensity by 14 ± 1, 35 ± 1, 50 ± 2, and 100 ± 3%, respectively (Fig. 7E). Microtubule asters were visible at 1- and 5-min intervals, with full reassembly observed by 30 min (Fig. 7D). Similar increases in microtubule intensities were observed in scrambled control cells, with no significant differences from the control at any time point (Fig. S7F). However, CEP41-depleted cells exhibited delayed recovery compared to control cells, with microtubule intensity increasing by 15 ± 0.6, 27 ± 0.5, 47 ± 2, and 62 ± 1% after 1, 5, 10, and 30 min, respectively. Recovery of CEP41-depleted cells at 5, 10, and 30 min was significantly lower than control cells, indicating a reduced rate of microtubule assembly. Specifically, the extent of microtubule nucleation seen at 5 min in control and scrambled control cells was not seen in CEP41-depleted cells, with a difference of 25% compared to control (Fig. 7E). Moreover, while no significant difference in microtubule intensity was observed between cells before nocodazole treatment and after 30 min of recovery in control and scrambled control cells, there was a 38% difference between the microtubule intensity of CEP41-depleted cells before nocodazole treatment and after 30 min recovery, suggesting incomplete microtubule reassembly possibly due to reduced nucleation rates (Fig. S7F). The results underscore the importance of CEP41 in maintaining the interphase microtubule network, which in turn is crucial for preserving cell shape.

### CEP41 is essential for cell proliferation and cell cycle progression

During the immunofluorescence analysis of CEP41-depleted cells, we observed that the total number of cells per field was lesser than the control and scrambled control. To investigate this further, we studied the effect of CEP41 depletion on cell proliferation. After 48 hours of transfection with shRNAs, cells were allowed to grow for one cycle. CEP41-depleted cells did not double in number in comparison to control untransfected cells. The percentage inhibition of cell proliferation was calculated to be 72 ± 17% for CEP41-depleted cells, whereas it was 20 ± 9% for scrambled control cells (Fig. 8A). Further, we checked the effect of CEP41 depletion on the cell cycle progression to determine if cell proliferation is inhibited due to a cell cycle block (Fig. 8B). CEP41-depleted and scrambled control cells were grown in puromycin-containing media and then subjected to flow cytometric analysis. Compared to control and scrambled control, there was a significant reduction in the number of CEP41-depleted cells in the G2/M phase. Compared to 29 ± 2% of control G2/M cells, 21 ± 2% of CEP41-depleted cells were in the G2/M phase. On the contrary, there was a slight increase in the number of cells in the S phase in the case of CEP41 depletion. We further subjected the cells to a double thymidine block to synchronize cells at the G1/S boundary (Fig. 8B). Synchronized cells were released from the thymidine block and allowed to grow for 8 hours to progress into mitosis. Control untransfected and scrambled control cells progressed into mitosis with 45 ± 1% and 44 ± 1% cells in the G2/M stage, respectively (Fig. 8B,C). However, CEP41-depleted cells did not progress to mitosis as efficiently as control cells, with only 27 ± 2% of cells in the G2/M phase. Interestingly, 54.3 ± 0.5% of CEP41-depleted cells remained in the G1 state post-thymidine release, compared to 40 ± 2% of control cells (Fig. 8B,C). The results suggest that CEP41 depletion slows down the progression of cells into the G2/M phase.

**Fig 8.**
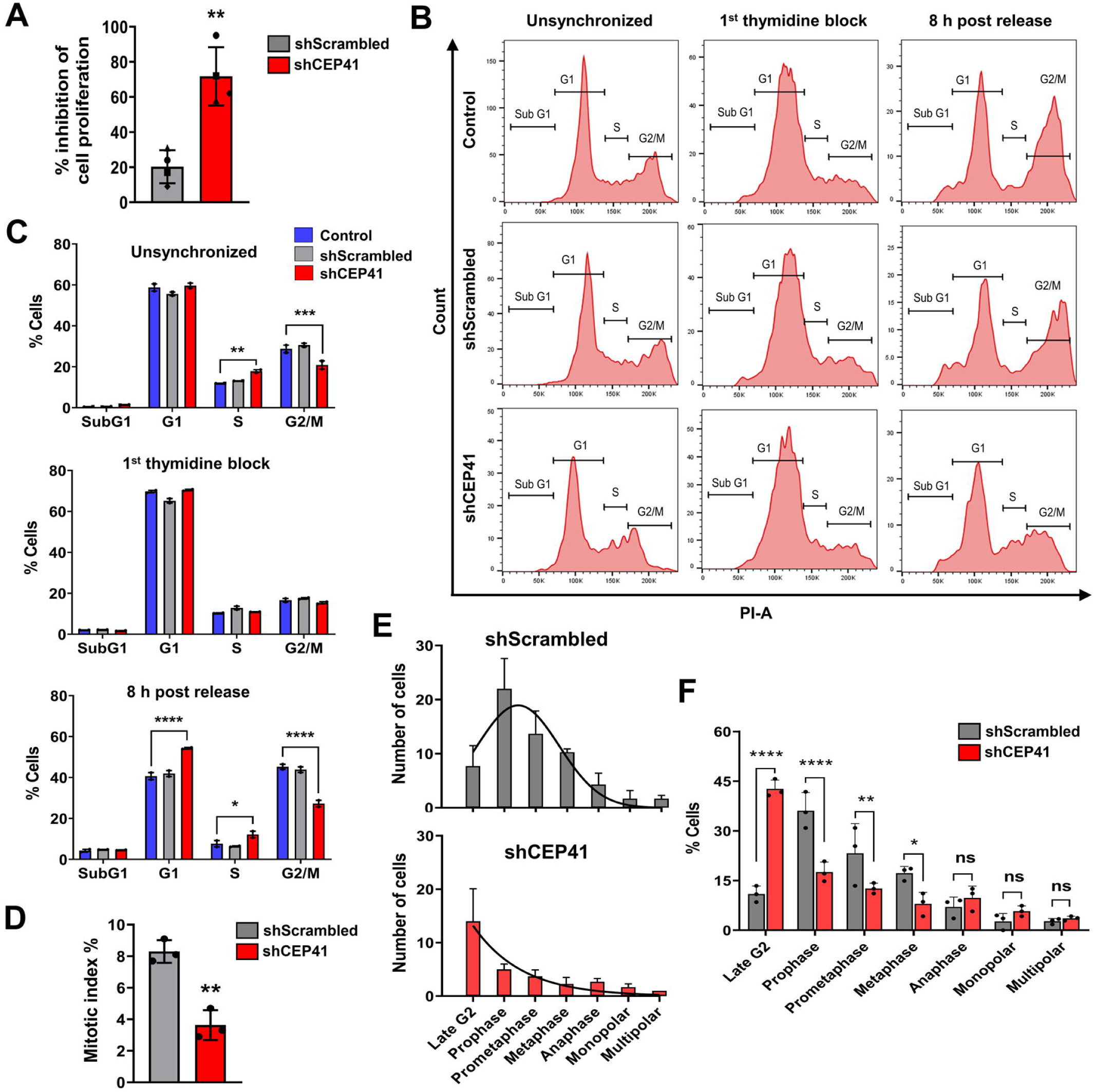
CEP41 depletion inhibited proliferation and cell cycle progression of NIH3T3 cells. **(A)** CEP41 depletion inhibited cell proliferation. NIH3T3 cells were transfected with scrambled or CEP41 shRNA. Percentage inhibition of cell proliferation was determined with respect to control. The data were an average of four independent sets represented as mean ± s.d. (\*\**P* < 0.01). Statistical significance was determined by Student’s *t*-test. **(B)** CEP41-depleted cells failed to progress to the G2/M phase. NIH3T3 cells transfected with scrambled or CEP41 shRNA were synchronized by a double thymidine block and, 8 h post-thymidine release, were stained with propidium iodide and subjected to flow cytometry analysis. Representative cytograms of cells unsynchronized, synchronized at the G1 stage (1^st^ thymidine block), and 8 h post thymidine release are shown. **(C)** Percentage of cells at different cell cycle stages. The data represent mean ± s.e.m. of two independent sets with a minimum of 2000 cells analyzed in each case (\**P* < 0.05; \*\**P* < 0.01; \*\*\**P* < 0.001; \*\*\*\**P* < 0.0001). Statistical significance was determined by two-way ANOVA test. **(D)** Mitotic index determination. NIH3T3 cells transfected with scrambled or CEP41 shRNA were fixed and stained for α-tubulin (red), phospho-Histone H3 (green), and DNA (blue). The mitotic index was determined as the ratio of the number of cells at mitosis / total number of cells x 100. Data represent the mean ± s.d. of three independent sets with 700 cells counted in each case (\*\**P* < 0.01). Statistical significance was determined by Student’s *t*-test. **(E)** Distribution of scrambled control and CEP41-depleted mitotic cells in different stages of mitosis. Data represent the mean ± s.d. of three independent sets, fitted in a non-linear regression curve. **(F)** Percentage of mitotic cells at different stages of mitosis. The data represent mean ± s.d. of three independent sets (ns *P* > 0.05; \**P* < 0.05; \*\*\**P* < 0.001; \*\*\*\**P* < 0.0001). Statistical significance was determined by two-way ANOVA test.

To further understand the effect of CEP41 depletion on mitotic progression, we determined the mitotic index for CEP41-depleted cells by calculating the ratio of the number of cells undergoing mitosis to the total number of cells (Fig. 8D). Cells were stained with phospho-histone H3 (pHH3) antibody, a mitotic marker, that recognizes the mitosis-specific phosphorylation of histone H3 (Ser10) (Hendzel et al., 1997). The mitotic index was estimated to be 8.3 ± 0.7% for scrambled control cells and 3.6 ± 0.9% for CEP41-depleted cells, representing a 57 ± 8% reduction in the number of mitotic cells compared to control, indicating fewer cells progressing into mitosis (Fig. 8D). Furthermore, based on DNA morphology and presence of spindle poles, we categorized pHH3-positive cells into different stages of mitosis (Fig. S8A,B). As the cell progresses from the G2 to M phase, nuclear DNA condenses into chromosomes, and duplicated centrosomes move apart to opposite ends to form spindle poles. Under the influence of dynamic microtubules with one end tethered to chromosomes, the chromosomes are aligned at the metaphase plate and then segregated equally towards spindle poles. Cells positive for pHH3 but lacking visible chromosomes and spindle poles were termed late G2 cells as these cells had condensed DNA, which is evident by the phosphorylated histone staining; however, they did not have distinct spindle poles (Fig. S8A,B). Cells with visible spindle poles but no chromosomes were termed prophase cells. Further cells with unaligned chromosomes were prometaphase, properly aligned chromosomes signified metaphase, and segregating chromosomes indicated anaphase. In scrambled control cells, the mitotic cells exhibited a normal distribution with cells distributed across late G2, prophase, prometaphase, metaphase, and anaphase (Fig. 8E). However, CEP41-depleted cells were notably more prevalent in the late G2 phase, constituting 43 ± 2% of mitotic cells compared to 11 ± 2% in control (Fig. 8E,F). Additionally, there was a proportional decrease in prophase, prometaphase, and metaphase cells in the case of CEP41 depletion, with respective counts of 18 ± 3, 13 ± 1, and 8 ± 3%, compared to 36 ± 5, 23 ± 9 and 17 ± 2% in control (Fig. 8E,F). The result suggests that CEP41 depletion slows down the progression of cells into mitosis, potentially by inhibiting mitotic entry. Overall, the results highlight the essential role of CEP41 in cell cycle regulation, possibly through its functions associated with the centrosome and the microtubules.

## Discussion

The key finding of our study is the identification of CEP41 as a microtubule-associated protein that promotes microtubule assembly and stabilization. We provide evidence that CEP41 is vital for maintaining the cellular microtubule network with indispensable roles in cell cycle progression and proliferation. We also show that the coiled-coil motifs and the rhodanese-like domain of CEP41 are essential for its interaction with the microtubules.

### CEP41 is a microtubule-interacting protein

CEP41 has been suggested to be a microtubule-interacting protein (Andersen et al., 2003; Gache et al., 2010); however, its direct binding and association with microtubules have not been shown. Here, we demonstrate CEP41’s binding to microtubules in cells and *in vitro*. In a co-sedimentation assay, purified CEP41 was sedimented with preformed microtubules, suggesting an apparent interaction between the proteins. Further, in prior studies, endogenous CEP41 was reported to localize only at the centrosome and ciliary axoneme. In our investigation, CEP41 colocalized with the microtubule lattice upon overexpression in HeLa and NIH3T3 cells. This observation resembles end-binding proteins like EB1 and CLIP-170, which, at endogenous concentrations, localize exclusively to growing microtubule tips yet bind throughout the microtubule lattice *in vitro* (Ligon et al., 2006). CEP41’s interaction with tubulin was also evident in co-immunoprecipitation and *in vitro* microtubule polymerization assays. Thus, the results suggest that, at endogenous levels, CEP41 could potentially be involved in microtubule anchoring and organization due to its direct binding to the lattice or soluble tubulin. Furthermore, CEP41 was detected in the MAP-enriched fraction of tubulin extracted from goat brain tissue, implying that it is a part of the MAP family of proteins, which are well known to bind to microtubules and regulate their assembly. MAPs can also modulate microtubule dynamics by influencing the interaction of other proteins with microtubules (Bodakuntla et al., 2019). CEP41 is involved in the recruitment of tubulin glutamylase TTLL6 to the ciliary axoneme; however, how it is involved in this regulation is unclear (Lee et al., 2012). We propose that it could be due to CEP41’s direct interaction with microtubules, where it could function as an intermediary and promote localization of TTLL6 to the ciliary microtubules. Similarly, cilia and spindle-associated protein (CSAP) have been reported to bind to TTLL5 and regulate its interaction with microtubules (Bompard et al., 2018). Another axoneme-associated protein, CEP162, is shown to mediate the interaction of transition zone components with ciliary microtubule (Wang et al., 2013). Establishing CEP41’s direct interaction with tubulin and microtubules provides valuable insights into its role in microtubule-dependent cellular functions. Furthermore, we demonstrated that the interaction between CEP41 and microtubules is mediated by its rhodanese-like domain and coiled-coil motifs. Coiled-coil motifs in centrosomal proteins are common, where they play an essential role in sub-cellular localization and contribute to protein-protein interactions (Salisbury, 2003). However, the rhodanese homology domain is primarily found in phosphatases and yeast ubiquitin-specific proteases (Bordo and Bork, 2002). Although catalytically inactive, the RHOD domain in CEP41 likely contributes to its stability and interactions with other proteins. Individuals with Joubert syndrome carry heterozygous mutations in the conserved coiled-coil region and the RHOD domain (Lee et al., 2012). When we examined these mutants, the R179H mutant exhibited impaired microtubule interaction. Located within the RHOD domain of CEP41, this mutation could impede CEP41’s ability to bind to microtubules, thereby disrupting its microtubule-associated functions and potentially contributing to disease pathology. This further underscores the crucial role of CEP41-microtubule interaction in cellular processes. Our assessment of the 3D structure of CEP41, generated by AlphaFold, also revealed that the R179 residue forms multiple polar contacts with neighboring residues, indicating its potential role in maintaining structural integrity. The impairment of microtubule binding resulting from a single mutation further highlights the intricate and specific nature of the interaction between CEP41 and microtubules. This suggests that subtle changes in the structure of CEP41 can significantly affect its ability to bind to microtubules effectively.

### CEP41 is a novel regulator of microtubule organization and bundling

Regulation of microtubule dynamics in cells is a complex process involving the coordinated activity of several proteins, including MAPs, centrosomal proteins, motor proteins, and several protein kinases that control the activation and recruitment of the nucleating factors (Bodakuntla et al., 2019; Kumari and Panda, 2018; Prassanawar and Panda, 2019; Tovey and Conduit, 2018). We propose CEP41 as another member of this regulatory network with newly identified microtubule-stabilizing activity. Upon CEP41 overexpression, we observed extensive microtubule bundling and an increase in the overall microtubule content in HeLa cells. Similarly, in CEP41-expressing NIH3T3 cells, a dense network of microtubules emanating from the centrosome was seen, suggesting that CEP41 positively regulates microtubule assembly. Further, purified CEP41 promoted *in vitro* microtubule assembly and prevented dilution-induced microtubule disassembly, again hinting at its function as a microtubule stabilizer. Microtubule nucleation is a cooperative process and requires the initial assembly of stable microtubule seeds that grow into polymers (Kuchnir Fygenson et al., 1995; Voter and Erickson, 1984). CEP41 might interact with tubulin and stabilize the initial oligomeric tubulin assemblies. TPX2, a microtubule-stabilizing protein, is thought to nucleate microtubules similarly by stabilizing oligomers (Schatz et al., 2003). The microtubule bundling observed in cells could also be due to an increase in the rate of microtubule assembly mediated by CEP41 overexpression. CEP41 directly interacts with microtubules; therefore, it can act as a cross-linker and enhance lateral interactions among the microtubules, prompting bundling. Another centrosomal and ciliary protein, ENKD1, has been shown to exhibit similar microtubule bundling and stabilization in cells and *in vitro* (Tiryaki et al., 2022). Moreover, in NIH3T3 cells, CEP41 depletion disrupted the interphase microtubule network and delayed microtubule reassembly, emphasizing its importance in organizing and maintaining the cellular microtubule array. Since CEP41 showed its capability to nucleate and stabilize microtubules, we propose its possible involvement in regulating microtubule nucleation at the centrosome. Although a direct interaction between CEP41 and γ-tubulin wasn’t detected in a co-immunoprecipitation assay (Fig. S8C), CEP41 may interact with other γ-TuRC components.

### CEP41 is essential for cell proliferation

During cell division, the centrosome orchestrates a complete reorganization of the interphase microtubule network into a dynamic mitotic spindle (Doxsey et al., 2005). This highly dynamic transition requires several centrosomal and microtubule-associated proteins that constantly modulate microtubule dynamics. Owing to the microtubule nucleating and stabilizing functions of CEP41, we hypothesized its involvement in the organization of spindle microtubules. Interestingly, CEP41 localized at the spindle poles and the midbody in dividing cells. In addition, CEP41 depletion inhibited cell proliferation and hampered cell cycle progression by delaying the transition of cells from the G1/S phase to the G2/M phase, validating our hypothesis that CEP41 is an essential protein with critical functions during cell division. An increase in the number of cells in the G1 phase also hints at faulty centrosome duplication, suggesting a possible role of CEP41 in centriole biogenesis or maturation, although this needs further assessment (Nigg, 2007). Earlier, the defective phenotypes of CEP41 knockdown were attributed only to ciliary dysfunctions (Ki et al., 2020; Lee et al., 2012). However, our findings suggest possible disruption of other cellular processes, specifically microtubule-dependent processes. For instance, retarded cell migration and impaired vascular development were prominent in CEP41-deficient zebrafish embryos (Ki et al., 2020), and our results suggest that it could also be due to CEP41’s role in microtubule organization. Similarly, CEP41 was reported to be essential for ciliogenesis and cilia disassembly (Graser et al., 2007; Lee et al., 2012), and here, we report its involvement in cell cycle regulation. Cilia disassembly and the onset of the cell cycle are tightly coordinated processes, with many centrosomal and ciliary proteins as key players functioning in this crosstalk (Izawa et al., 2015; Jackson, 2011). IFT88 is one example of an essential ciliary protein that is reported to regulate the G1/S transition in non-ciliated cells (Robert et al., 2007). Similarly, CEP164, a centrosomal protein, regulates the G2/M transition and is vital for primary cilia formation (Graser et al., 2007). CEP41 could also be a key player in this crosstalk that warrants further investigation. The presence of a conserved RHOD domain in CEP41, the function of which remains unexplored, strongly hints at the possibility of CEP41’s involvement in cell cycle regulation. The RHOD domain is found in the catalytic domain of Cdc25 phosphatases and the non-catalytic domain of MAPK-phosphatases (Bordo and Bork, 2002). Both of these classes of proteins play roles in regulating various signaling pathways associated with cell growth and survival. Interestingly, CEP41 was shown to be essential for angiogenesis through its role in activating Aurora kinase A and HIF1α (hypoxia-inducible factor 1α), which regulates the cilia disassembly pathway (Ki et al., 2020). Therefore, investigating CEP41’s role in such regulatory pathways will be an exciting endeavor.

Overall, the results identify a potential role of CEP41 as a centrosomal MAP involved in maintaining stable microtubule structures in cells and provide a background to explore its functions as a cell cycle regulator.

## Materials and methods

### Construction of expression vectors

Full-length cDNA encoding human CEP41 (GenBank accession no. NM_018718.3) cloned in pcDNA3.1 vector with a C-terminal eGFP tag was procured from Origene (USA). The CEP41 cDNA was amplified and subcloned into a bacterial expression vector pET28a(+) at Nde1 and SalI restriction sites with an N-terminal hexahistidine tag. Deletion constructs with deleted N-terminal (ΔN: residues deleted 33-137), rhodanese-like domain (ΔRHOD: 169-266), C-terminal (ΔC: 273-373), and both N- and C- terminal (ΔNC: 33-137 and 273-373) regions were generated by PCR using full-length CEP41 in pcDNA3.1 as a template. CEP41-GFP constructs with disease-causing mutations M36T, Q89E, R179H (Lee et al., 2012), and P206A (Patowary et al., 2019) were generated by site-directed mutagenesis using PCR with full-length CEP41 in pcDNA3.1 as a template. All constructs were purified using endotoxin-free plasmid extraction kits (Qiagen) and sequenced to verify intact reading frames.

### Cell culture and transfection

Mouse embryo-derived fibroblast (NIH3T3) and human cervical carcinoma (HeLa) cells were obtained from the cell repository of the National Centre for Cell Sciences (India). The cells were cultured in Dulbecco’s Modified Eagle’s Medium (HiMedia) supplied with 10% (v/v) fetal bovine serum (Invitrogen) and 1% (v/v) antibiotic-antimycotic solution (HiMedia). The cells were free from microbial or fungal contamination and were maintained at 37°C in a humidified cell culture incubator (Sanyo) with 5% CO_2_. Cells at 60-70% confluency were transiently transfected with plasmids or shRNAs using Lipofectamine 3000 according to the manufacturer’s protocol (Invitrogen).

### Cell lysis and immunoblotting

Immunoblotting analysis was performed as described earlier (Kumari et al., 2021; Srivastava and Panda, 2018). Briefly, cells were trypsinized, lysed in a cold lysis buffer, and centrifuged, and the protein concentration of the resulting supernatant was determined using Bradford’s reagent (Bradford, 1976). Equal concentrations of the samples were resolved on SDS-PAGE gel and electroblotted onto PVDF membranes (Merck Millipore, USA) (Srivastava and Panda, 2018). The blots were blocked at room temperature (25°C) for 1 h with 5% skim milk in TBST (Tris-buffered saline pH 7.5 with 0.1% Tween-20), followed by incubation with primary antibody at 4°C overnight and peroxidase-conjugated secondary antibody at 25°C for 1 h. The blots were developed using the chemiluminescence of luminol (Thermo Fisher Scientific), transferred on X-ray film, and band intensities were quantified using ImageJ (National Institutes of Health, Bethesda, USA). Images of uncropped blots are shown in Fig. S9.

Primary antibodies used were mouse anti-α-tubulin (T9026, Sigma-Aldrich, at 1:1000), rabbit anti-CEP41 (17566-1-AP, Proteintech, at 1:800), mouse anti-β-actin (A2228, Sigma-Aldrich, at 1:2000), mouse anti-GAPDH (sc-47724, Santa Cruz Biotech, at 1:500), mouse anti-polyhistidine (H1029, Sigma-Aldrich, at 1:1000), mouse anti-EB1 (sc-47704, Santa Cruz Biotech, at 1:500), rabbit anti-Histone H3 (H0164, Sigma-Aldrich, at 1:5000), mouse anti-polyglutamylated tubulin (T9822, Sigma-Aldrich, at 1:1000), and mouse anti-γ-tubulin (T6557, Sigma-Aldrich, at 1:1000). Secondary antibodies used were horseradish peroxidase (HRP)-conjugated anti-mouse IgG (7076S, Cell Signaling Technology) and anti-rabbit IgG (1706615, Bio-Rad) at 1:2000.

### Co-immunoprecipitation assay

NIH3T3 cells seeded in T-75 flasks were harvested at 80-90% confluency and lysed in a cold lysis buffer (50 mM Tris, 250 mM NaCl, 5 mM EDTA, 50 mM NaF, 0.1% Triton X, 500 μM sodium orthovanadate, 2 mM PMSF, PIC, pH 7.4). The lysate was subjected to co-immunoprecipitation with desired antibodies (Kumari et al., 2021). The cell lysate was pre-cleared with Protein A agarose beads (Millipore, Sigma), and 700 μg of the pre-cleared lysate was incubated with 2 μg of either anti-CEP41, anti-α-tubulin, rabbit IgG, or mouse IgG at 4°C on a rocker overnight, followed by incubation with pre-washed Protein A agarose beads for 2 h. The beads were collected by centrifugation, washed, and the resin-bound immune complex was resuspended in an SDS loading buffer and boiled for 5 min. The samples were analyzed using SDS-PAGE followed by western blotting. Rabbit IgG, mouse IgG, and Protein A agarose beads incubated with pre-cleared lysate, without any antibody, were used as negative controls. The experiment was performed twice.

### Protein expression and purification

Recombinant full-length CEP41 was synthesized in *E.coli* BL21 (DE3) pLysE cells using the pET vector expression system with induction conditions of 30°C for 8 h with 0.5 mM IPTG. Post-induction, the cells were collected by centrifugation, lysed, and sonicated in ice-cold lysis buffer (50 mM Tris, 300 mM NaCl, 1mg/ml lysozyme, 2 mM PMSF, 0.1% Triton X-100, 0.01% β-ME, 10% glycerol, pH 8). Subsequently, the clarified cell lysate was incubated with pre-equilibrated Ni-NTA resin (Takara) for 2 h at 4°C in slow shaking conditions. The his-tagged protein was eluted with an elution buffer (50 mM Tris, 300 mM NaCl, and 250 mM imidazole, pH 8), concentrated, and dialyzed in a storage buffer (25 mM Tris, 50 mM NaCl, 10% glycerol, and pH 7). The dialyzed protein was aliquoted, snap-frozen, and stored at −80°C. Before every experiment, the histidine tag was cleaved using Thrombin CleanCleave Kit (Thermo Fisher Scientific), per the manufacturer’s protocol. The thrombin-bound beads were separated by centrifugation, and the supernatant passed through the Ni-NTA column. The resulting cleaved CEP41 was concentrated and analyzed by SDS-PAGE and immunoblotted using anti-CEP41 and anti-polyhistidine antibodies.

Tubulin was purified from the goat brain tissue, as described previously (Hamel and Lin, 1981; Panda et al., 2005). Briefly, the goat brain tissue was homogenized in PEM buffer (50 mM PIPES, 3 mM MgCl_2_, 1 mM EGTA, pH 6.8) containing 2 mM PMSF and 0.1% β-Me and centrifuged at 33,000 g at 4°C. The clear cell extract was polymerized at 37°C with 1 M glutamate, 10% DMSO, and 0.5 mM GTP (Sigma-Aldrich), centrifuged, and then depolymerized on ice. Two cycles of such polymerization and depolymerization were performed, and the resulting pure tubulin was snap-frozen and stored at −80°C. The purity of the protein was assessed by SDS-PAGE analysis.

Similarly, microtubule-associated proteins were isolated from the goat brain tissue employing two cycles of microtubule assembly and disassembly, as described earlier (Panda et al., 2005). After centrifugation, the clarified brain tissue extract was polymerized with 4 M glycerol and 0.5 mM GTP, followed by centrifugation and cold depolymerization. The presence of CEP41 in the MAP-rich fractions was detected by resolving 150 μg of the brain tissue extract and supernatant from the 1^st^ and 2^nd^ depolymerization fractions on SDS-PAGE gel and immunoblotted with anti-CEP41, anti-EB1, and anti-Histone H3 antibody. The experiment was performed twice.

### Microtubule binding assay

The binding of CEP41 with microtubules was determined, as described earlier (Asthana et al., 2012). Tubulin (30 μM) was polymerized with taxol (20 μM) and 1 mM GTP in PEM buffer. 3 μM of the polymerized microtubules were incubated with CEP41 (1, 2, 3, and 5 μM) for 20 min at 37°C in PEM buffer containing taxol (5 μM) and 1 mM GTP. The reaction mixture was centrifuged at 60,000 g for 20 min at 30°C in an ultracentrifuge Optima MAX-XP (Beckman Coulter), and the resulting microtubule pellet was gently washed with warm PEM buffer containing 10% DMSO. The pellet was dissolved in SDS running buffer, and equal volumes of the soluble and polymer fractions were resolved on 12% SDS-PAGE gel and immunoblotted with anti-CEP41 antibody. As a control, CEP41 (1, 2, 3, and 5 μM), without microtubules, was incubated and centrifuged in the same conditions and analyzed by SDS-PAGE.

### Light scattering assay

The effect of CEP41 on *in vitro* microtubule assembly was studied, as mentioned before (Chaudhary et al., 2016). Tubulin (30 μM) was incubated with CEP41 (1, 2, 3, and 5 μM) for 20 min on ice in PEM buffer, followed by the addition of GTP (1 mM). The reaction kinetics was monitored at 37°C by measuring light scattering at 350 nm using Spectramax M2e (Molecular Devices). Bovine serum albumin (BSA) and alkaline phosphatase (10 μM) were used as negative controls. Similarly, the effect of CEP41 on taxol-induced tubulin polymerization was determined. Tubulin (14 μM) was incubated with CEP41 (1, 2, 3, and 5 μM), followed by the addition of taxol (4 μM) and GTP (1 mM), and the reaction kinetics were measured. The rate of tubulin polymerization was calculated from the slope of the linear part (elongation phase) of the polymerization curve.

### Sedimentation assay

Tubulin (30 μM) was polymerized with CEP41 (1, 2, 3, and 5 μM) in PEM buffer and GTP (1 mM) at 37°C for 30 min. The reaction mixture was ultracentrifuged at 50,000 g at 30°C for 30 min. The supernatant was separated, and the pellet was washed with warm PEM buffer with 10% DMSO. The pellet was dissolved in SDS running buffer, and equal volumes of supernatant and pellet were resolved on 15% SDS-PAGE. Similarly, the effect of CEP41 on taxol-induced tubulin polymerization was determined. Tubulin (14 μM) was incubated with CEP41 (1, 2, 3, and 5 μM) for 20 min on ice in PEM buffer, followed by the addition of taxol (4 μM) and GTP (1 mM). The reaction mixture was polymerized for 30 min at 37°C, followed by centrifugation and SDS-PAGE analysis.

### Transmission electron microscopy

Tubulin (30 μM) was polymerized in the presence of CEP41 (5 μM) and GTP (1 mM) in PEM buffer for 30 min at 37°C, and the microtubules formed were imaged in an electron microscope as described before (Asthana et al., 2012). Briefly, the polymerized mixture was spotted on carbon-formvar coated copper grids, air-dried, washed with double distilled water, and stained with 0.75% uranyl acetate. The grids were imaged in an electron microscope, JEM 2100 ultra HRTEM instrument at 200 kV (JEOL). The experiment was performed three times.

### Dilution-induced disassembly assay

The effect of CEP41 on dilution-induced microtubule depolymerization was determined, as described earlier (Asthana et al., 2012). Tubulin (20 μM) was polymerized with 10% DMSO and 1 mM GTP in PEM buffer. The polymerized microtubules were diluted 10-fold in PEM buffer containing 1 mM GTP and CEP41 (1, 3, and 5 μM) and incubated for 10 min at 37°C. The reaction mixture was then ultracentrifuged at 60,000 g for 20 min at 30°C. The supernatant was separated, and the pellet was gently washed with warm PEM buffer containing 10% DMSO. The pellet was dissolved in SDS running buffer, and equal volumes of the soluble and polymer fractions were resolved on 12% SDS-PAGE gel, followed by immunoblotting using anti-α-tubulin and anti-CEP41 antibodies.

### Immunofluorescence and microscopy

Cells were seeded at 50,000 cells/ml density on clean glass coverslips in 24-well plates and incubated overnight. After incubation, cells were transfected with desired plasmids and incubated for 24 h, followed by fixation and immunostaining, as described earlier (Asthana et al., 2013; Kumari and Panda, 2022). Briefly, the cells were fixed with 3.7% formaldehyde, followed by permeabilization with 0.1% Triton-X 100 and blocking with 4% BSA, and then incubated with primary antibodies in BSA for 3 h at 25°C, followed by incubation with secondary antibodies for 1 h at 25°C. DNA was stained with Hoechst (10 μg/ml) for 15 min at 25°C, and following three washes, the coverslips were mounted and cured overnight. Primary antibodies used were mouse anti-α-tubulin (T9026, Sigma-Aldrich), rabbit anti-acetyl-α-tubulin (Lys40) (SAB5600134, Sigma-Aldrich), mouse anti-polyglutamylated tubulin (T9822, Sigma-Aldrich), rabbit anti-phospho-Histone H3 (pSer^10^) (H0412, Sigma-Aldrich), and mouse anti-γ-tubulin (T6557, Sigma-Aldrich) at 1:400. Secondary antibodies were Alexa Fluor 594-conjugated anti-mouse IgG (A11005, Invitrogen) and anti-rabbit IgG (A11012, Invitrogen), and FITC-conjugated anti-mouse IgG (F0257, Sigma-Aldrich) and anti-rabbit IgG (F0382, Sigma-Aldrich), at 1:400. Actin was stained using Alexa Fluor 488-conjugated phalloidin following the manufacturer’s protocol (A12379, Invitrogen). Images were acquired in z-stacks in a spinning-disc CSU-X1 (Yokogawa) or a laser scanning LSM 780 (Zeiss) confocal microscope using a 63x oil immersion lens. The exposure and gain settings were kept identical among the control and treated groups to quantify fluorescence intensities. Images were processed in Zen Blue software, and intensities were quantified using ImageJ. Colocalization analyses of GFP-expressing cells were done using the Coloc 2 plugin available in ImageJ Fiji software. Measurement of cell area was done in Zen Blue software by drawing cell boundaries based on microtubule staining. All experiments were performed thrice, with a minimum of 50 cells quantified for each group.

For the nocodazole-induced disassembly assay, HeLa cells transiently transfected with CEP41-GFP and GFP constructs were incubated for 24 h and then treated with 500 nM nocodazole (Sigma-Aldrich) for 1 h and subjected to immunostaining as described above. For imaging CEP41-expressing mitotic cells, HeLa cells were transfected with CEP41-GFP and GFP constructs and then synchronized by double thymidine block using 2 mM thymidine (Sigma-Aldrich) (Ma and Poon, 2011). Post 8 h of thymidine release, cells were fixed and immunostained.

### *In vitro* microtubule staining

The microtubules formed in the presence of CEP41 were visualized by immunostaining (Asthana et al., 2012; Girotra et al., 2017). Tubulin (30 μM) was polymerized with CEP41 (5 μM) and GTP (1 mM) in PEM buffer for 30 min at 37°C, and the polymerized solution was loaded on clean poly-L-lysine coated coverslips, incubated for 20 min, and blot dried. The coverslips were fixed with 3.7% formaldehyde and immunostained using α-tubulin antibody. The experiment was performed three times.

### CEP41 depletion using shRNA

The endogenous level of CEP41 in NIH3T3 cells was depleted using shRNA, as described before (Kumari et al., 2023). Cells were seeded in 60 mm dishes at 50-60% confluency and transiently transfected with CEP41 shRNA-1 (5’-CCCTACCAGCTGAGAATAAAT-3’, TRCN0000142809, Sigma-Aldrich), CEP41 shRNA-2 (5’-GCTTACAGTTACCCAATTGCA-3’, TRCN0000143499, Sigma-Aldrich), or scrambled shRNA (SHC002, Sigma-Aldrich) using Lipofectamine 3000. Post-transfection, the cells were incubated for 48 h, harvested, and lysed. The levels of CEP41 were determined by western blotting using anti-CEP41 and anti-β-actin (loading control) antibodies. For immunostaining of depleted cells, NIH3T3 cells were grown on coverslips in 24-well plates, transfected with CEP41 and scrambled shRNA, and 48 h post-transfection subjected to immunostaining as described above.

For the microtubule reassembly assay, NIH3T3 cells were cultured on coverslips in 24-well plates. Subsequently, they were transfected with CEP41 and scrambled shRNA, and 48 h post-transfection, treated with 500 nM nocodazole for 1 h. Following the treatment, the nocodazole was removed by washing cells with cold media three times. The cells were then replenished with warm media and allowed to recover for 1, 5, 10, and 30 min at 37°C. At each designated time point, the soluble fraction of cells was extracted by incubating the cells with extraction media (PEM + 0.1% Triton X-100) for 45 sec at 37°C. Subsequently, the cells were rapidly fixed with 100% chilled methanol and subjected to immunostaining following the procedure described above. Images were acquired in z-stacks in a spinning-disc confocal microscope using a 63x oil immersion lens. The exposure and gain settings were kept identical among the control and treated groups to quantify fluorescence intensities. Images were processed in Zen Blue software, and intensities were quantified using ImageJ. The cell boundary was determined based on the brightfield images. The experiment was performed twice, with 100 cells quantified for each experimental group.

### Cell proliferation assay

Sulforhodamine B (SRB) assay was performed to determine the effect of CEP41 depletion on cell proliferation. NIH3T3 cells were seeded in 96-well plates (8000 cells/well) and transfected with CEP41 and scrambled shRNA, as mentioned above. After 48 h, the cells were fixed with chilled 50% trichloroacetic acid and processed for the SRB assay as described before (Ray et al., 2007; Skehan et al., 1990). The percentage inhibition of cell proliferation was determined, and the experiment was performed four times.

### Cell cycle analysis

The effect of CEP41 depletion on cell cycle progression was determined by flow cytometric analysis (Gajula et al., 2013; Mohan and Panda, 2008). CEP41-depleted and scrambled control cells were grown in puromycin (4 μg/ml)-containing medium for 2 weeks to enrich shRNA-expressing cells. The cells were then synchronized by a double thymidine block and 8 h post-thymidine release, trypsinized, fixed with 70% ethanol, and stained with propidium iodide (30 μg/ml). Using a BD FACSAria flow cytometer (Becton Dickinson), a minimum of 2000 cells were analyzed in each group, and the experiment was performed twice. The data were processed using the FlowJo software (Tree Star).

### Determination of mitotic index

The effect of CEP41 depletion on mitotic progression was determined by estimating the mitotic index as described earlier (Srivastava and Panda, 2018). NIH3T3 cells were cultured on coverslips in 24-well plates, transfected with CEP41 and scrambled shRNA, and 48 h post-transfection, subjected to immunostaining as described above. The cells were stained with α-tubulin and phospho-histone H3 (pHH3) antibody. DNA was stained with Hoechst. The cells were visualized under an Eclipse TE 2000U microscope (Nikon, Japan) at 63x magnification. Mitotic index was calculated as the ratio of the number of cells undergoing mitosis to the total number of cells. The experiment was performed thrice, and 700 cells were counted in each case. Representative images were acquired in z-stacks in a spinning-disc confocal microscope.

### In silico analysis

The secondary and tertiary structures of CEP41 were predicted using PSI-PRED (McGuffin et al., 2000) and AlphaFold (Jumper et al., 2021; Varadi et al., 2022), respectively. The image was created using PyMoL (DeLano, 2002). Coiled-coil regions in CEP41 were predicted using Ncoils in the Waggawagga prediction software (Simm et al., 2015). Sequence alignment of CEP41 orthologues was carried out using Clustal Omega (Sievers et al., 2011).

### Statistics

Data were analyzed and plotted using GraphPad Prism version 8.0.2 and represented as mean ± s.d or mean ± s.e.m. with error bars denoting s.d or s.e.m. Experiments were performed as three independent sets unless mentioned otherwise. An unpaired, two-tailed *t*-test was used to determine statistical significance between the mean of two groups, and for significance between three or more groups, a one-way or two-way analysis of variance (ANOVA) was applied, followed by Tukey’s multiple-comparison test. *P* < 0.05 was considered statistically significant.

## Acknowledgments

The authors thank IIT Bombay for the central facilities of spinning disk and laser scanning confocal microscopes, FACS, and cryo-HRTEM. We thank Prof. Subba Rao Gangi Shetty (IISc, Bangalore) for providing the shRNA clones of CEP41. S.S.P. thanks the Council of Scientific and Industrial Research, Government of India, for her doctoral fellowship.

## Data availability

All relevant data can be found within the article and its supplementary information.

## References

Andersen, J. S., Wilkinson, C. J., Mayor, T., Mortensen, P., Nigg, E. A. and Mann, M. (2003). Proteomic characterization of the human centrosome by protein correlation profiling. Nature 426, 570–574.

Asthana, J., Kuchibhatla, A., Jana, S. C., Ray, K. and Panda, D. (2012). Dynein light chain 1 (LC8) association enhances microtubule stability and promotes microtubule bundling. Journal of Biological Chemistry 287, 40796–40805.

Asthana, J., Kapoor, S., Mohan, R. and Panda, D. (2013). Inhibition of HDAC6 deacetylase activity increases its binding with microtubules and suppresses microtubule dynamic instability in MCF-7 cells. Journal of Biological Chemistry 288, 22516–22526.

Bettencourt-Dias, M., Hildebrandt, F., Pellman, D., Woods, G. and Godinho, S. A. (2011). Centrosomes and cilia in human disease. Trends in Genetics 27, 307–315.

Bodakuntla, S., Jijumon, A. S., Villablanca, C., Gonzalez-Billault, C. and Janke, C. (2019). Microtubule-Associated Proteins: Structuring the Cytoskeleton. Trends Cell Biol 29, 804–819.

Bompard, G., van Dijk, J., Cau, J., Lannay, Y., Marcellin, G., Lawera, A., van der Laan, S. and Rogowski, K. (2018). CSAP Acts as a Regulator of TTLL-Mediated Microtubule Glutamylation. Cell Rep 25, 2866–2877.

Bordo, D. and Bork, P. (2002). The rhodanese/Cdc25 phosphatase superfamily. Sequence-structure-function relations. EMBO Rep 3, 741–746.

Bradford, M. (1976). A Rapid and Sensitive Method for the Quantitation of Microgram Quantities of Protein Utilizing the Principle of Protein-Dye Binding. Anal Biochem 72, 248–254.

Bre, M. H. and Karsenti, E. (1990). Effects of brain microtubule-associated proteins on microtubule dynamics and the nucleating activity of centrosomes. Cell Motil Cytoskeleton 15, 88–98.

Chaudhary, V., Venghateri, J. B., Dhaked, H. P. S., Bhoyar, A. S., Guchhait, S. K. and Panda, D. (2016). Novel Combretastatin-2-aminoimidazole Analogues as Potent Tubulin Assembly Inhibitors: Exploration of Unique Pharmacophoric Impact of Bridging Skeleton and Aryl Moiety. J Med Chem 59, 3439–3451.

Conduit, P. T., Wainman, A. and Raff, J. W. (2015). Centrosome function and assembly in animal cells. Nat Rev Mol Cell Biol 16, 611–624.

DeLano, W. L. (2002). The PyMOL Molecular Graphics System, Version 1.1. Schrodinger LLC.

Doxsey, S. (2001). Re-evaluating centrosome function. Nat Rev Mol Cell Biol 2, 688–698.

Doxsey, S., Zimmerman, W. and Mikule, K. (2005). Centrosome control of the cell cycle. Trends Cell Biol 15, 303–311.

Gache, V., Waridel, P., Winter, C., Juhem, A., Schroeder, M., Shevchenko, A. and Popov, A. V. (2010). Xenopus meiotic microtubule-associated interactome. PLoS One 5, e9248.

Gajula, P. K., Asthana, J., Panda, D. and Chakraborty, T. K. (2013). A synthetic dolastatin 10 analogue suppresses microtubule dynamics, inhibits cell proliferation, and induces apoptotic cell death. J Med Chem 56, 2235–2245.

Girotra, M., Srivastava, S., Kulkarni, A., Barbora, A., Bobra, K., Ghosal, D., Devan, P., Aher, A., Jain, A., Panda, D., et al. (2017). The C-terminal tails of heterotrimeric kinesin-2 motor subunits directly bind to α-tubulin1: Possible implications for cilia-specific tubulin entry. Traffic 18, 123–133.

Graser, S., Stierhof, Y. D., Lavoie, S. B., Gassner, O. S., Lamla, S., Le Clech, M. and Nigg, E. A. (2007). Cep164, a novel centriole appendage protein required for primary cilium formation. Journal of Cell Biology 179, 321–330.

Hamel, E. and Lin, C. M. (1981). Glutamate-Induced Polymerization of Tubulin - Characteristics of the Reaction and Application to the Large-Scale Purification of Tubulin. Arch Biochem Biophys 209, 29–40.

Izawa, I., Goto, H., Kasahara, K. and Inagaki, M. (2015). Current topics of functional links between primary cilia and cell cycle. Cilia 4, 1–13.

Jackson, P. K. (2011). Do cilia put brakes on the cell cycle? Nat Cell Biol 13, 340–342.

Jakobsen, L., Vanselow, K., Skogs, M., Toyoda, Y., Lundberg, E., Poser, I., Falkenby, L. G., Bennetzen, M., Westendorf, J., Nigg, E. A., et al. (2011). Novel asymmetrically localizing components of human centrosomes identified by complementary proteomics methods. EMBO Journal 30,.

Jumper, J., Evans, R., Pritzel, A., Green, T., Figurnov, M., Ronneberger, O., Tunyasuvunakool, K., Bates, R., Žídek, A., Potapenko, A., et al. (2021). Highly accurate protein structure prediction with AlphaFold. Nature 596, 1520–1535.

Karr, T. L. and Purich, D. L. (1979). A microtubule assembly/disassembly model based on drug effects and depolymerization kinetics after rapid dilution. Journal of Biological Chemistry 254, 10885–10888.

Ki, S. M., Kim, J. H., Won, S. Y., Oh, S. J., Lee, I. Y., Bae, Y., Chung, K. W., Choi, B., Park, B., Choi, E., et al. (2020). CEP41-mediated ciliary tubulin glutamylation drives angiogenesis through AURKA -dependent deciliation. EMBO Rep 21, e48290.

Korvatska, O., Estes, A., Munson, J., Dawson, G., Bekris, L. M., Kohen, R., Yu, C. E., Schellenberg, G. D. and Raskind, W. H. (2011). Mutations in the TSGA14 gene in families with autism spectrum disorders. *American Journal of Medical Genetics*, Part B: Neuropsychiatric Genetics 156, 303–311.

Kuchnir Fygenson, D., Flyvbjerg, H., Sneppen, K., Libchaber, A. and Leibler, S. (1995). Spontaneous nucleation of microtubules. Phys Rev E 51, 5058.

Kumar, N. (1981). Taxol-induced polymerization of purified tubulin. Mechanism of action. Journal of Biological Chemistry 256, 10435–10441.

Kumari, A. and Panda, D. (2018). Regulation of microtubule stability by centrosomal proteins. IUBMB Life 70, 602–611.

Kumari, A. and Panda, D. (2022). Monitoring the Disruptive Effects of Tubulin-Binding Agents on Cellular Microtubules. In Microtubules: Methods and Protocols (ed. H. Inaba), pp. 431-448. Humana, New York, NY: Springer.

Kumari, A., Shriwas, O., Sisodiya, S., Santra, M. K., Guchhait, S. K., Dash, R. and Panda, D. (2021). Microtubule-targeting agents impair kinesin-2-dependent nuclear transport of β-catenin: Evidence of inhibition of Wnt/β-catenin signaling as an important antitumor mechanism of microtubule-targeting agents. FASEB Journal 35, e21539.

Kumari, A., Prassanawar, S. S. and Panda, D. (2023). β-III Tubulin Levels Determine the Neurotoxicity Induced by Colchicine-Site Binding Agent Indibulin. ACS Chem Neurosci 14, 19–34.

Lee, J. E., Silhavy, J. L., Zaki, M. S., Schroth, J., Bielas, S. L., Marsh, S. E., Olvera, J., Brancati, F., Iannicelli, M., Ikegami, K., et al. (2012). CEP41 is mutated in Joubert syndrome and is required for tubulin glutamylation at the cilium. Nat Genet 44, 193–199.

Ligon, L. A., Shelly, S. S., Tokito, M. K. and Holzbaur, E. L. F. (2006). Microtubule binding proteins CLIP-170, EB1, and p150Glued form distinct plus-end complexes. FEBS Lett 580, 1327–1332.

Ma, H. T. and Poon, R. Y. C. (2011). Synchronization of HeLa Cells. In Methods in Molecular Biology (ed. G. Banfalvi), pp 151-161. Humana, New York, NY: Springer

McGuffin, L. J., Bryson, K. and Jones, D. T. (2000). The PSIPRED protein structure prediction server. Bioinformatics 16, 404–405.

Mohan, R. and Panda, D. (2008). Kinetic stabilization of microtubule dynamics by estramustine is associated with tubulin acetylation, spindle abnormalities, and mitotic arrest. Cancer Res 68, 6181–6189.

Nigg, E. A. (2007). Centrosome duplication: of rules and licenses. Trends Cell Biol 17, 215–221.

Nigg, E. A. and Raff, J. W. (2009). Centrioles, Centrosomes, and Cilia in Health and Disease. Cell 139, 663–678.

Panda, D., Rathinasamy, K., Santra, M. K. and Wilson, L. (2005). Kinetic suppression of microtubule dynamic instability by griseofulvin: implications for its possible use in the treatment of cancer. Proceedings of the National Academy of Sciences 102, 9878–9883.

Patowary, A., Won, S. Y., Oh, S. J., Nesbitt, R. R., Archer, M., Nickerson, D., Raskind, W. H., Bernier, R., Lee, J. E. and Brkanac, Z. (2019). Family-based exome sequencing and case-control analysis implicate CEP41 as an ASD gene. Transl Psychiatry 9, 4.

Piperno, G., LeDizet, M. and Chang, X. J. (1987). Microtubules containing acetylated alpha-tubulin in mammalian cells in culture. J Cell Biol 104, 289–302.

Prassanawar, S. S. and Panda, D. (2019). Tubulin heterogeneity regulates functions and dynamics of microtubules and plays a role in the development of drug resistance in cancer. Biochemical Journal 476, 1359–1376.

Ray, S., Mohan, R., Singh, J. K., Samantaray, M. K., Shaikh, M. M., Panda, D. and Ghosh, P. (2007). Anticancer and antimicrobial metallopharmaceutical agents based on palladium, gold, and silver N-heterocyclic carbene complexes. J Am Chem Soc 129, 15042–15053.

Reiter, J. F. and Leroux, M. R. (2017). Genes and molecular pathways underpinning ciliopathies. Nat Rev Mol Cell Biol 18, 533–547.

Robert, A., Margall-Ducos, G., Guidotti, J. E., Brégerie, O., Celati, C., Bréchot, C. and Desdouets, C. (2007). The intraflagellar transport component IFT88/polaris is a centrosomal protein regulating G1-S transition in non-ciliated cells. J Cell Sci 120, 628–637.

Roostalu, J. and Surrey, T. (2017). Microtubule nucleation: Beyond the template. Nat Rev Mol Cell Biol 18, 702–710.

Salisbury, J. L. (2003). Centrosomes: Coiled-coils organize the cell center. Current Biology 13, R88–R90.

Schatz, C. A., Santarella, R., Hoenger, A., Karsenti, E., Mattaj, I. W., Gruss, O. J. and Carazo-Salas, R. E. (2003). Importin α-regulated nucleation of microtubules by TPX2. EMBO Journal 22,.

Sievers, F., Wilm, A., Dineen, D., Gibson, T. J., Karplus, K., Li, W., Lopez, R., McWilliam, H., Remmert, M., Söding, J., et al. (2011). Fast, scalable generation of high-quality protein multiple sequence alignments using Clustal Omega. Mol Syst Biol 7, 2060–2070.

Simm, D., Hatje, K. and Kollmar, M. (2015). Waggawagga: Comparative visualization of coiled-coil predictions and detection of stable single α-helices (SAH domains). Bioinformatics 31, 767–769.

Skehan, P., Storeng, R., Scudiero, D., Monks, A., Mcmahon, J., Vistica, D., Warren, J. T., Bokesch, H., Kenney, S. and Boyd, M. R. (1990). New colorimetric cytotoxicity assay for anticancer-drug screening. J Natl Cancer Inst 82, 1107–1112.

Srivastava, S. and Panda, D. (2018). A centrosomal protein STARD9 promotes microtubule stability and regulates spindle microtubule dynamics. Cell Cycle 17, 2052–2068.

Tiryaki, F., Deretic, J. and Firat-Karalar, E. N. (2022). ENKD1 is a centrosomal and ciliary microtubule-associated protein important for primary cilium content regulation. FEBS Journal 289, 3789–3812.

Tovey, C. A. and Conduit, P. T. (2018). Microtubule nucleation by γ-tubulin complexes and beyond. Essays Biochem 62, 765–780.

Vallee, R. B. (1986). Reversible assembly purification of microtubules without assembly-promoting agents and further purification of tubulin, microtubule-associated proteins, and MAP fragments. Methods Enzymol 134, 89–104.

Varadi, M., Anyango, S., Deshpande, M., Nair, S., Natassia, C., Yordanova, G., Yuan, D., Stroe, O., Wood, G., Laydon, A., et al. (2022). AlphaFold Protein Structure Database: Massively expanding the structural coverage of protein-sequence space with high-accuracy models. Nucleic Acids Res 50, D439–D444.

Vertii, A., Bright, A., Delaval, B., Hehnly, H. and Doxsey, S. (2015). New frontiers: discovering cilia-independent functions of cilia proteins. EMBO Rep 16, 1275–1287.

Voter, W. A. and Erickson, H. P. (1984). The kinetics of microtubule assembly. Evidence for a two-stage nucleation mechanism. Journal of Biological Chemistry 259, 10430–10438.

Wang, W. J., Tay, H. G., Soni, R., Perumal, G. S., Goll, M. G., MacAluso, F. P., Asara, J. M., Amack, J. D. and Bryan Tsou, M. F. (2013). CEP162 is an axoneme-recognition protein promoting ciliary transition zone assembly at the cilia base. Nat Cell Biol 15, 591–601.

