## Supplementary information for "CEP41, a ciliopathy-linked centrosomal protein, regulates microtubule assembly and cell proliferation"

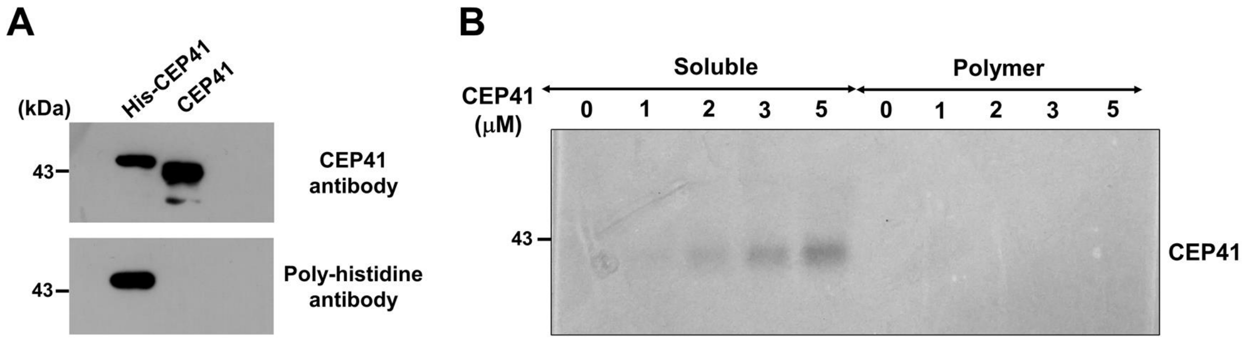


**Fig S1. CEP41 does not sediment and remains in the soluble fraction. (A)** Representative western blot to confirm the purification of his-tagged CEP41 and cleavage of the histidine tag. **(B)** Representative SDS-PAGE gel image showing sedimentation of indicated concentrations of CEP41 when subjected to ultracentrifugation in the absence of microtubules.

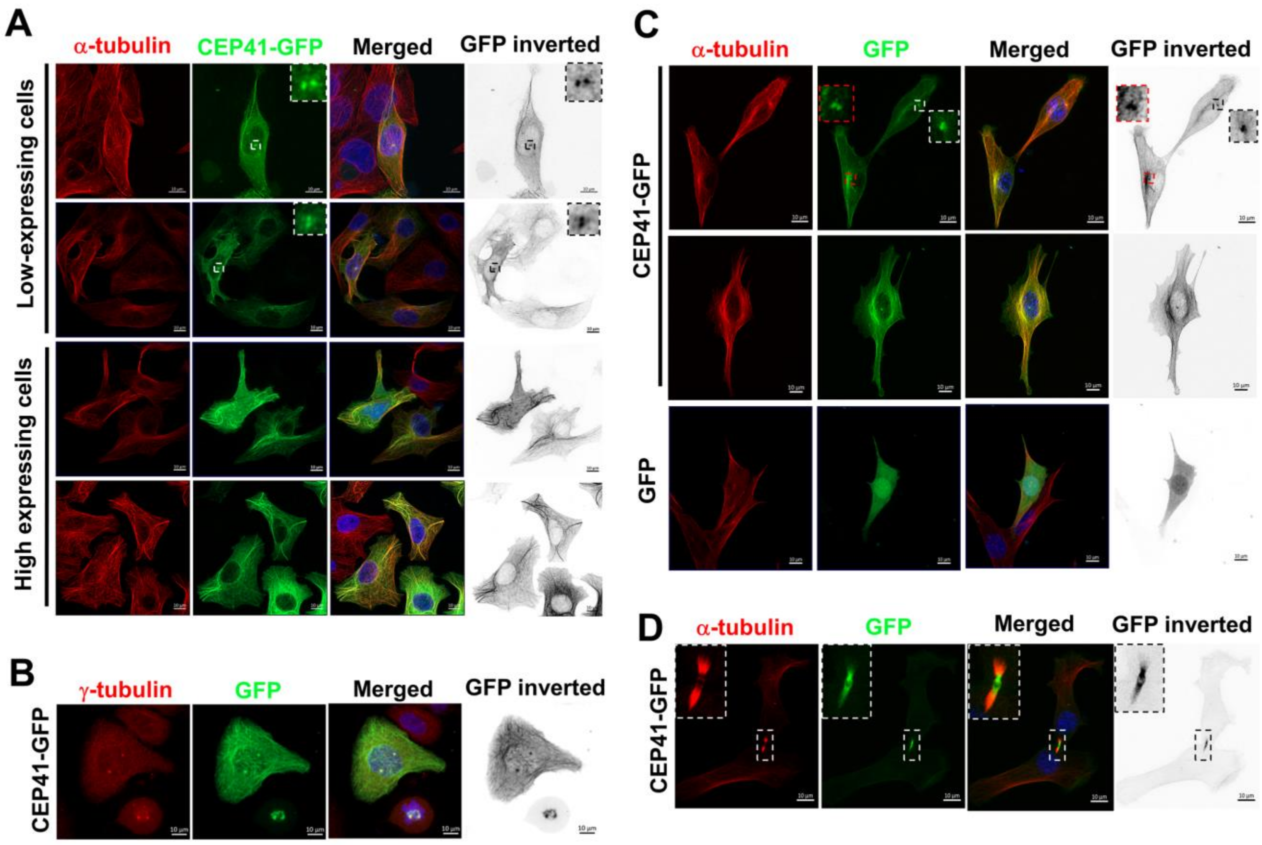


**Fig S2. Localization of CEP41-GFP in HeLa and NIH3T3 cells.** HeLa cells transfected with CEP41-GFP or GFP constructs were immunostained for **(A)** α-tubulin (red) or **(B)** γ-tubulin (red), and DNA was stained with Hoechst (blue). Scale bar: 10 µm. **(A)** CEP41 depicts centrosome-like localization in low-expressing cells (top two panels) and localizes to microtubules in high-expressing cells (bottom two panels). **(B)** CEP41 colocalizes with γ-tubulin in interphase and mitotic cells. **(C)** NIH3T3 cells transfected with CEP41-GFP or GFP constructs and immunostained for α-tubulin (red), and DNA was stained with Hoechst (blue). Scale bar: 10 µm. CEP41 localizes to the centrosomes and microtubules in interphase cells **(C)** and at the midbody during cytokinesis **(D)**.

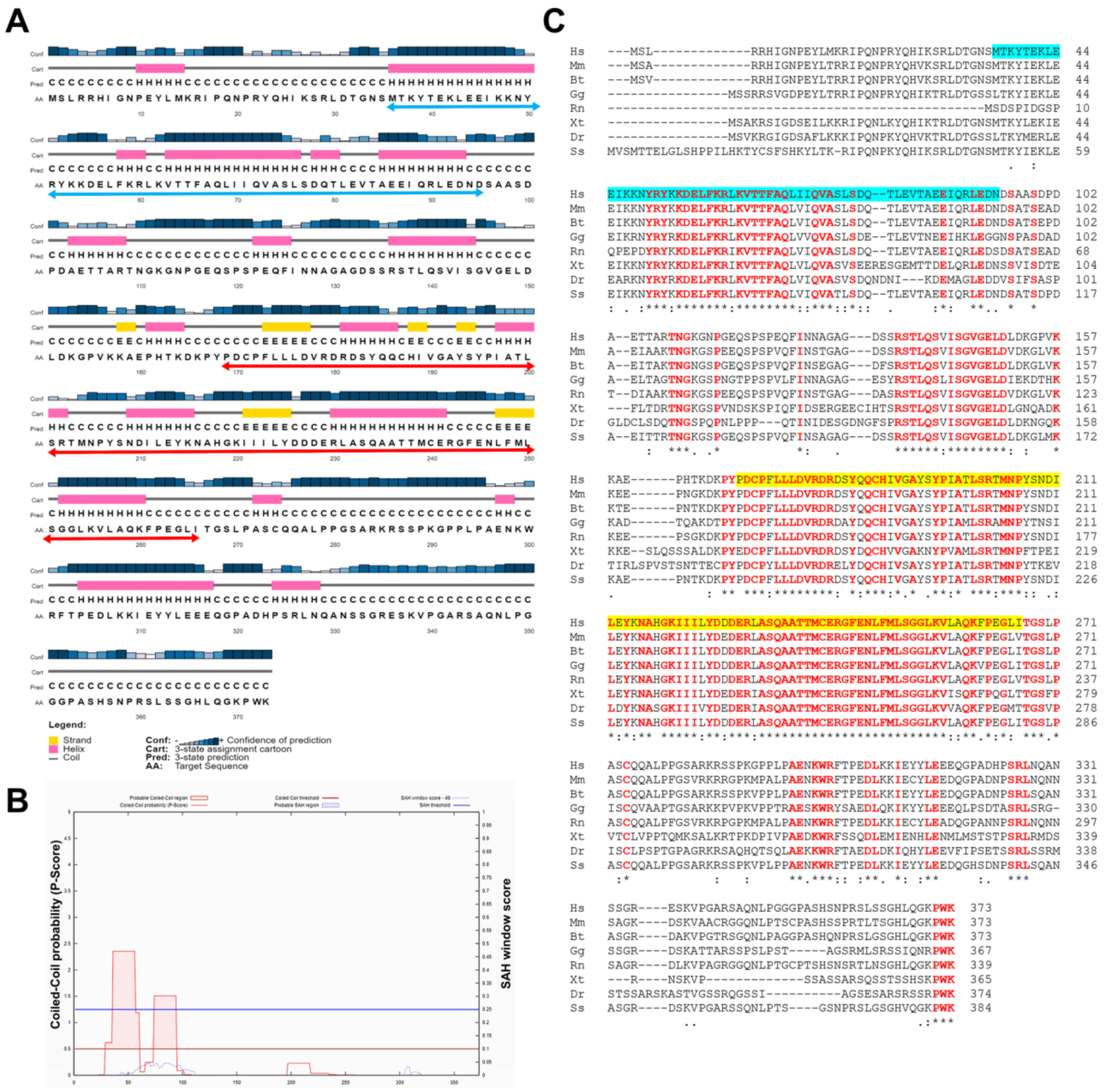


**Fig S3. Structure prediction of CEP41. (A)** Secondary structure prediction of CEP41 by PSI-PRED, arrows denote coiled-coil motif (cyan) and RHOD domain (red). **(B)** Prediction of coiled-coil regions in CEP41 using Ncoils in the Waggawagga prediction software. **(C)** Multiple sequence alignment of CEP41 and its orthologues. Sequence alignment was carried out using Clustal Omega. Hs – *Homo sapiens*, Mm – *Mus musculus*, Bt – *Bos taurus*, Gg – *Gallus gallus*, Rn – *Rattus norvegicus*, Xt – *Xenopus tropicalis*, Dr – *Danio rerio*, Ss – *Sus scrofa*. Coiled-coil motifs are highlighted in cyan, and the RHOD domain is yellow. The conserved residues are highlighted in red color.

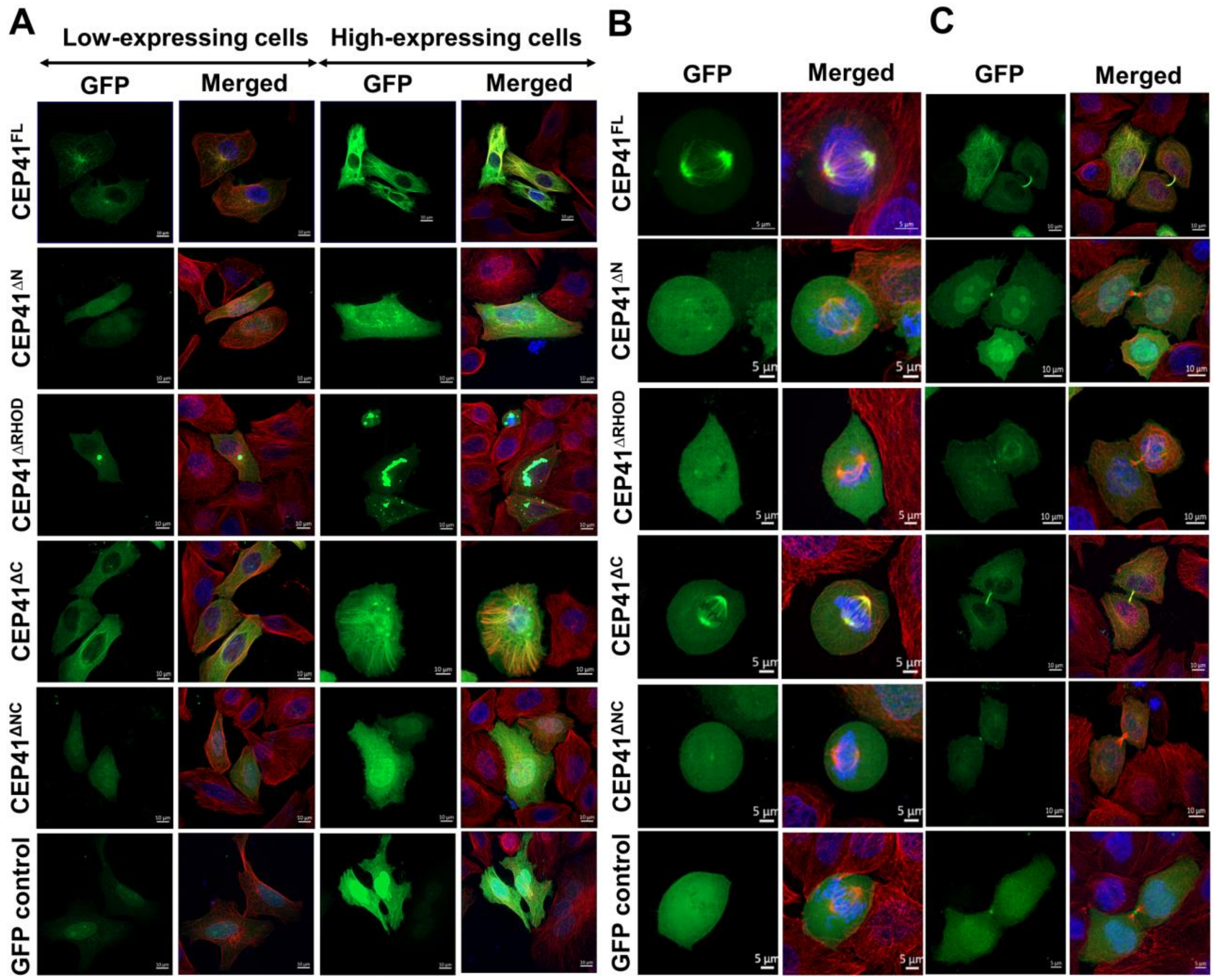


**Fig S4. Localization of deletion constructs of CEP41 in HeLa cells.** Fluorescence microscopy images of HeLa cells transfected with GFP-tagged full-length and truncated CEP41 constructs (green). Transfected cells were immunostained for $\alpha$-tubulin (red) 24 h post-transfection. DNA was stained with Hoechst (blue). Images were captured in a spinning-disk confocal microscope at 63x magnification. Localization of truncated mutants of CEP41 in low and high-expressing cells **(A),** at the spindle poles **(B),** and the midbody **(C)**. The scale bar is 5 µm **(B)** and 10 µm **(A and C)**.

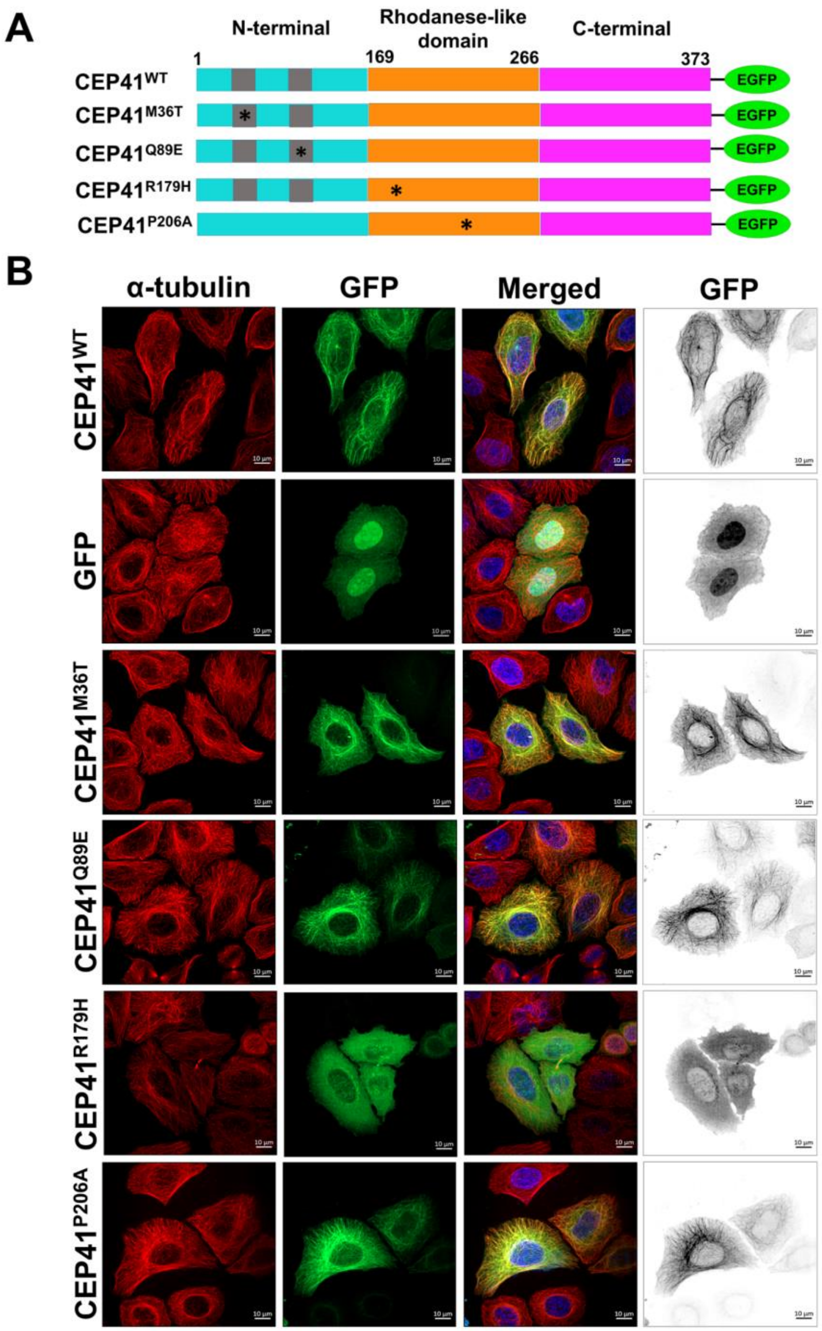


**Fig S5. Effect of disease-causing mutations on CEP41's interaction with microtubules. (A)** Mutated constructs of CEP41 with disease-causing mutations. Grey boxes indicate the predicted coiled-coil motifs, and asterisks indicate the position of the point mutations. **(B)** Fluorescence microscopy images of HeLa cells transfected with GFP-tagged full-length and mutated CEP41 constructs (green). Transfected cells were immunostained for $\alpha$-tubulin (red) 24 h post-transfection. DNA was stained with Hoechst (blue). Images were captured in a spinning-disk confocal microscope at 63x magnification. Scale bar: 10 $\mu$m.

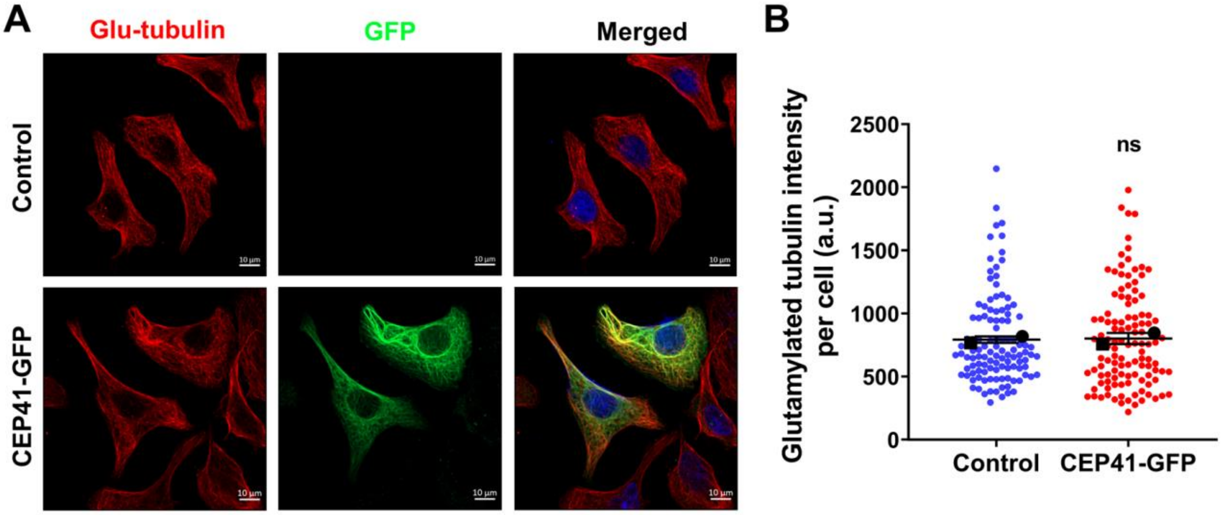


**Fig S6. Overexpression of CEP41-GFP does not change glutamylated tubulin levels in HeLa cells.** (A) Fluorescence microscopy images of HeLa cells transfected with CEP41-GFP or GFP constructs and immunostained for polyglutamylated tubulin (red). DNA was stained with Hoechst (blue). Scale bar: 10 μm. **(B)** The intensity of polyglutamylated tubulin per cell was quantified for 100 cells in each case. Data represent the mean ± s.e.m. of two independent sets (ns $P > 0.05$). Statistical significance was determined by Student's *t*-test.

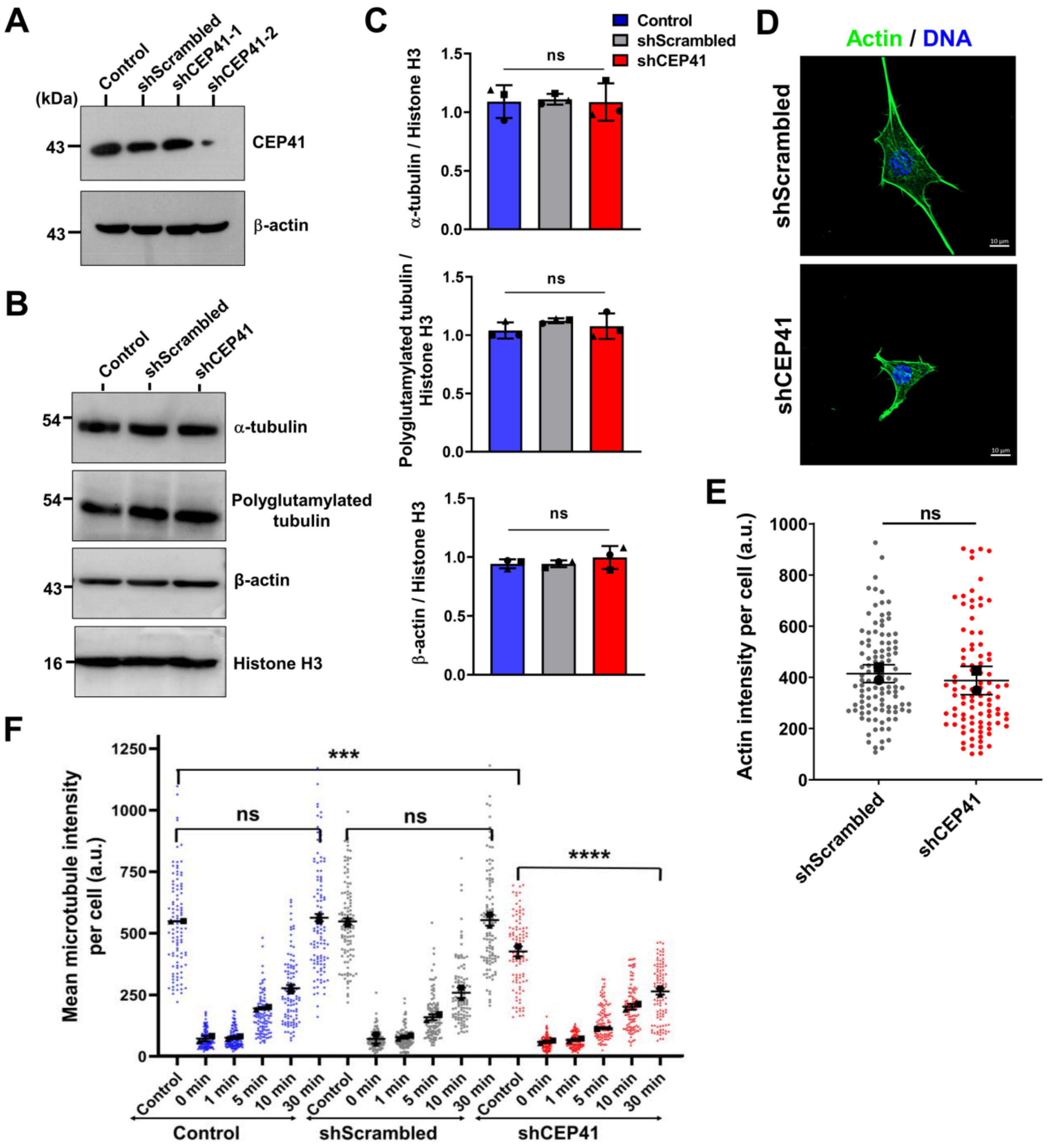


**Fig S7. CEP41 depletion in NIH3T3 cells. (A)** Representative immunoblot showing CEP41 levels in NIH3T3 cells transfected with scrambled or CEP41 shRNAs. β-actin was used as a loading control. **(B)** Representative immunoblots showing levels of indicated proteins in NIH3T3 cells transfected with scrambled or CEP41 shRNA. Histone H3 was used as a loading control. **(C)** Band intensities were quantified using ImageJ, and intensity ratios were plotted (ns $P > 0.05$). Error bars represent the standard deviation from three independent sets of experiments. **(D)** Fluorescence microscopy images of NIH3T3 cells transfected with scrambled or CEP41 shRNA and immunostained for actin with Alexa Fluor 488-

conjugated phalloidin (green). DNA was stained with Hoechst (blue). Scale bar: 10 μm. **(E)** The intensity of actin per cell was quantified for 100 cells in each case. Data represent the mean ± s.e.m. of two independent sets (ns $P > 0.05$). Statistical significance was determined by Student's *t*-test. **(F)** CEP41 depletion reduced the extent of microtubule reassembly. NIH3T3 cells transfected with scrambled or CEP41 shRNA were treated with nocodazole (500 nM) for 1 h to disassemble interphase microtubules. Post-treatment, nocodazole was washed out, and microtubule reassembly kinetics were monitored by incubating cells at 37°C. At indicated time points, the soluble fraction was extracted, and cells were fixed and immunostained. Microtubule intensity of 100 cells in each case were quantified using ImageJ (ns $P > 0.05$; ***$P < 0.001$; ****$P < 0.0001$). There was no significant difference between control and scrambled control intensities at any time point. Data represent the mean ± s.e.m. of two independent experiments. Statistical significance was determined by one-way ANOVA test.

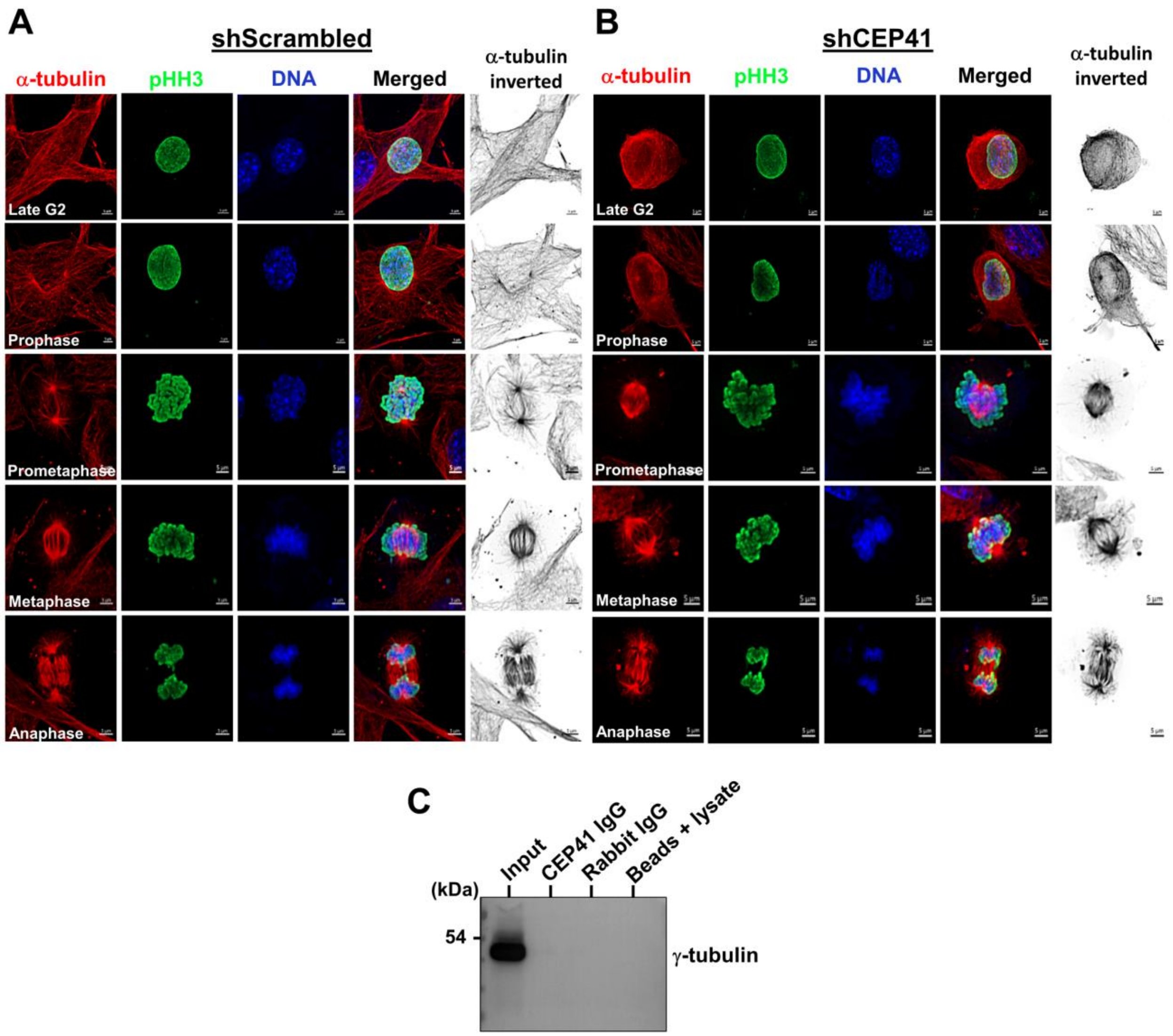


**Fig S8**. **Effect of CEP41 depletion on mitotic progression.** NIH3T3 cells transfected with scrambled or CEP41 shRNA were fixed and stained for α-tubulin (red), phospho-Histone H3 (green), and DNA (blue). Images were captured in a spinning disk confocal microscope at 63x magnification. Scale bar: 5 µm. Representative images of scrambled control **(A)** and CEP41-depleted **(B)** cells in different stages of mitosis. **(C)** CEP41 does not interact with γ-tubulin. Co-immunoprecipitation of CEP41 and γ-tubulin from NIH3T3 cells. The whole cell lysate was incubated with CEP41 IgG and immunoprecipitated using Protein A agarose beads. The input and eluates were immunoblotted with γ-tubulin antibody. The experiment was performed twice, and a representative blot is shown. Rabbit IgG and beads + lysate are negative controls.

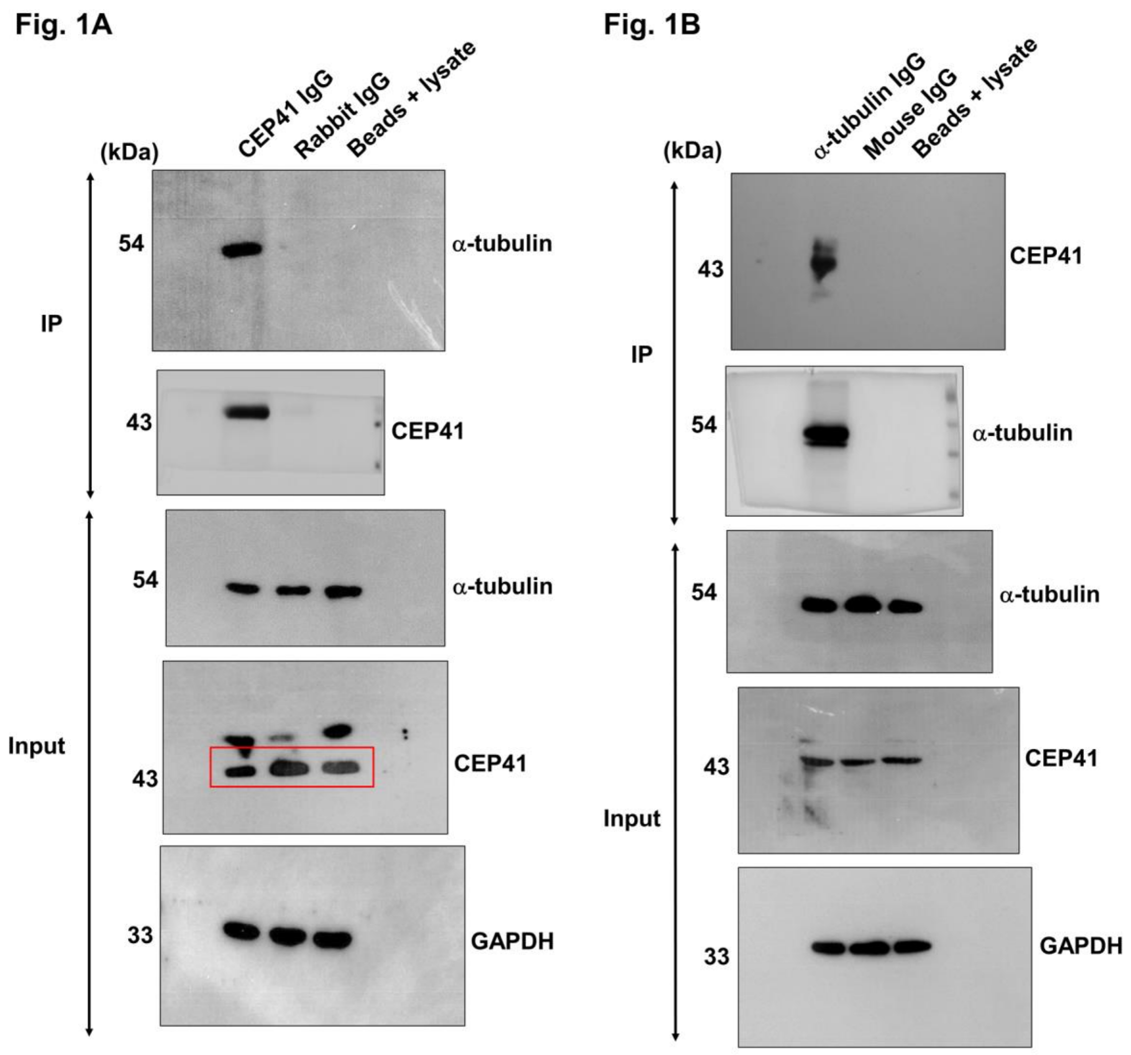


**Fig. S9. Blot Transparency.** Images of the uncropped blots and SDS gels for all corresponding figures and supplementary figures.

Fig. 1C

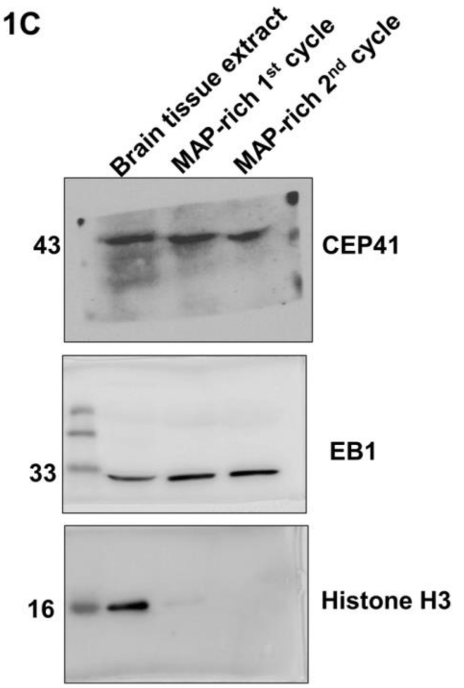


Fig. 1D

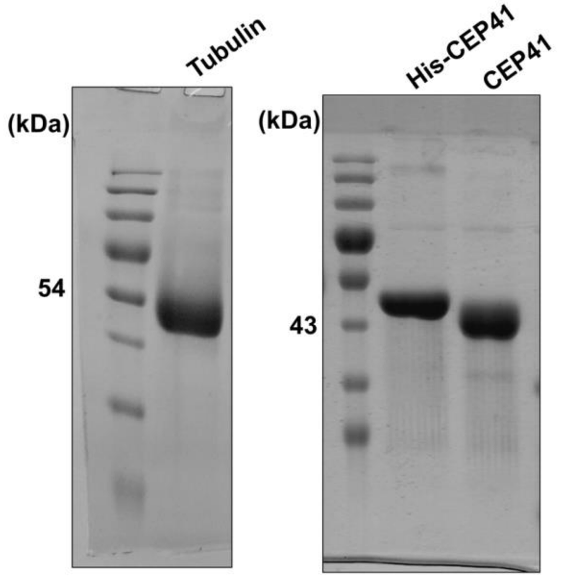


Fig. 1E

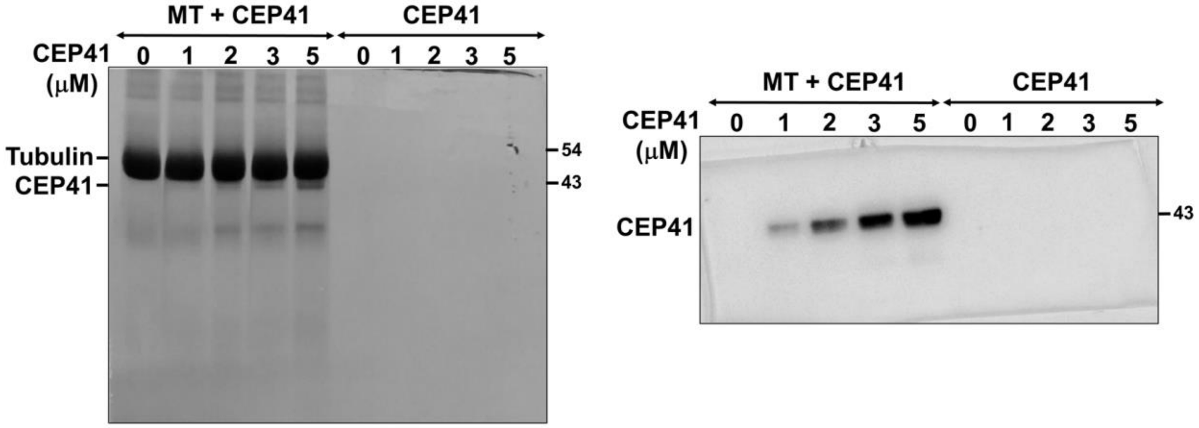

**Fig. 2C**

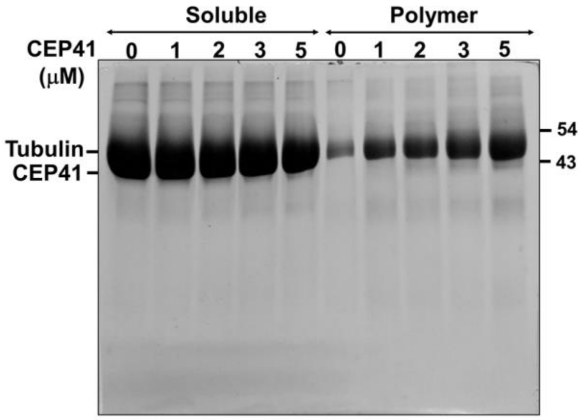


**Fig. 3C**

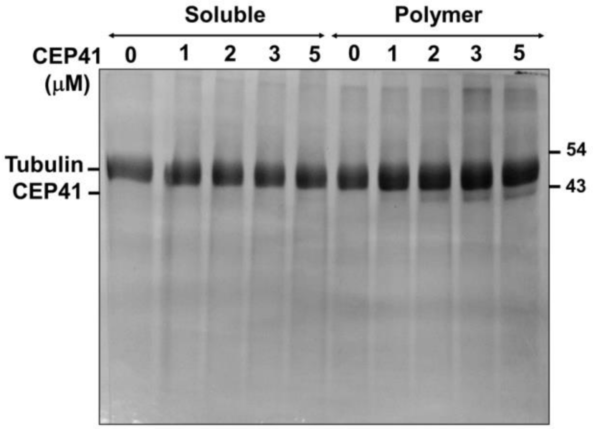


**Fig. 3E**

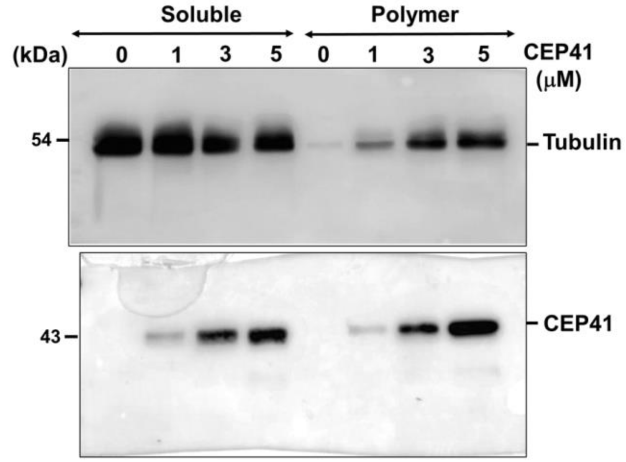


**Fig. 7A**

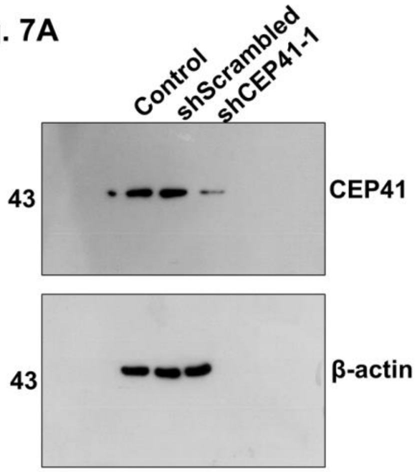

Fig. S1A

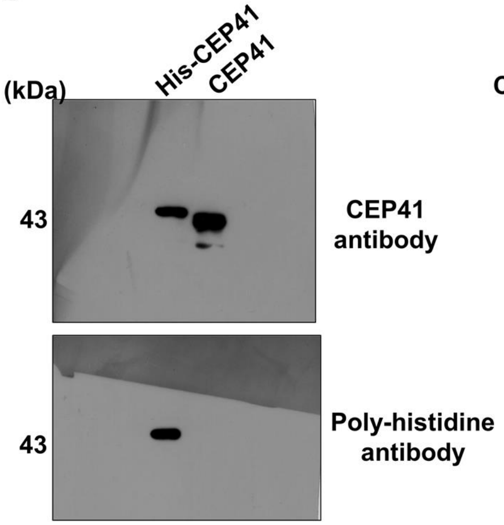


Fig. S1B

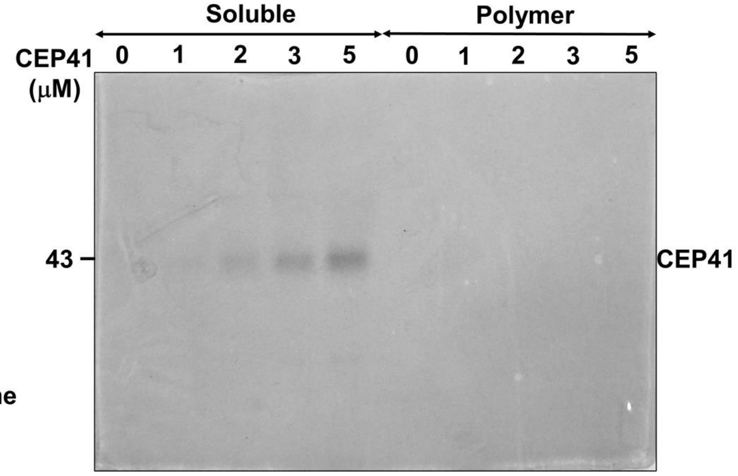


Fig. S7A

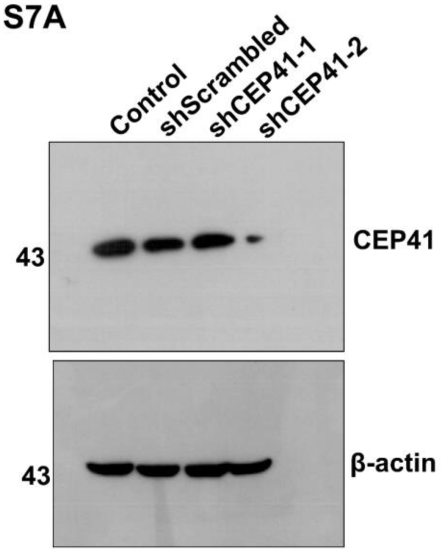


Fig. S7B

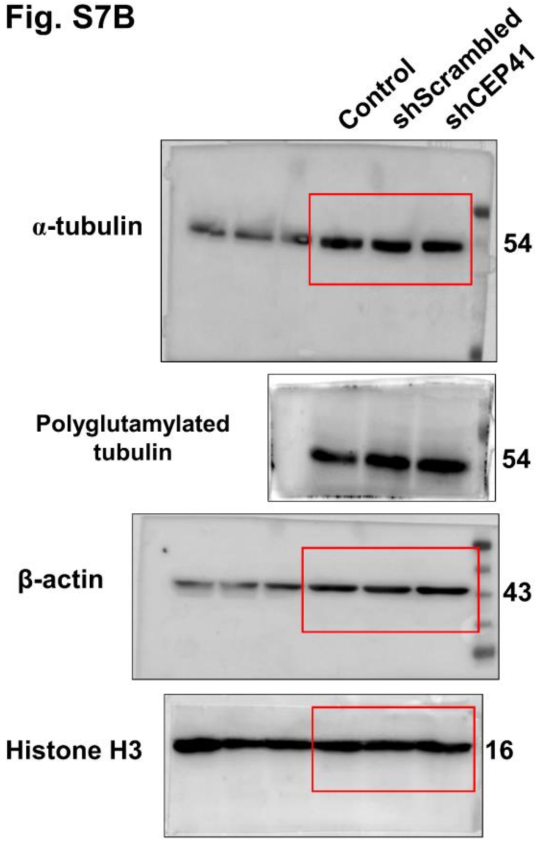


Fig. S8C

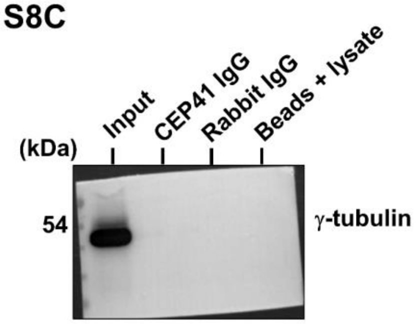
